# Bimodal Mechanical Response of Membrane Necks: Implications for the Nuclear Envelope

**DOI:** 10.1101/2025.03.10.642015

**Authors:** Beatrice J. Geiger, Weria Pezeshkian

## Abstract

Among the fascinating shapes that biomembranes exhibit are stomatocytes with multiple membrane necks, found for example in nuclear membranes and open autophagosomes. These morphologies, characterised by a high topological genus, can be visualised as spherical double membranes connected by neck-like structures. The necks are often occupied by specific biomolecular complexes, such as the nuclear pore complex, which divide the space into three distinct compartments. Understanding how the size of these necks responds to pressure gradients is fundamentally important for unravelling the influence of mechanical stimuli on traffic control through the necks, for example, in nuclear mechanosensing. In this work, we use computer simulations and theoretical analysis to investigate how neck size responds to variations in pressure or tension. Our findings reveal a two-phase behaviour: below a certain threshold, necks constrict as the pressure gradient increases, while above that threshold, they dilate. This response stems from the pure membrane’s mechanics and depends on the magnitude of the pressure gradient, the initial diameter of the neck and the membrane bending rigidity. We also provide a simple equation that links the threshold tension, the neck diameter and the bending rigidity, offering a useful tool to quickly assess different scenarios. Our results furthermore show that protein complexes in the neck partially counteract both constriction and dilation, stabilising neck size while preserving the same two-phase response to membrane tension. These findings uncover a promising, previously overlooked membrane property with implications for organelle shape and function, as well as for synthetic membrane design.

## MAIN TEXT

Biological and synthetic membranes assume a variety of shapes that are crucial for cell functioning. An interesting example is the stomatocyte shape, a concave inward structure resembling a bowl, often with tight openings. From a biomedical point of view, stomatocytes can be an indication of disease related to red blood cells ^1^. Stomatocyte morphologies are commonly observed under certain physical conditions. For instance, a spherical vesicle with a large surface-to-volume ratio or high osmotic pressure can undergo transitions into a stomatocyte shape ^2,3^. This can also be achieved through negative spontaneous curvature ^4^, representing the effect of transbilayer asymmetry due to distinct molecular compositions of each membrane leaflet and differences in the environments they face ^5,6^. In membranes with high topological genera, stomatocytic structures can form with multiple openings (**Fig. 1b**, Supplementary Fig. 1a) ^7–10^. The number of openings equals the surface’s topological genus plus one (g+1). Once an opening narrows (from here on referred to as membrane necks) specific molecular machinery can assemble to render the neck impermeable to selected biomolecules, effectively dividing the space into three compartments (see also **Fig. 1a**) ^4,11^. Similar structures are observed in cellular membranes, such as autophagosomes and the nuclear envelope ^12–18^. For example, in case of the nuclear envelope, its high-genus surface can be described as two quasi-spherical membranes connected by multiple membrane necks. The nuclear pore complex effectively regulates biomolecular trafficking through the necks ^19–23^. Because of the many potential implications on organelle shape and cell function, it is therefore essential to understand how the size of these necks is regulated in response to common external stimuli and macroscopic parameters, such as osmotic pressure.

**Fig. 1:**
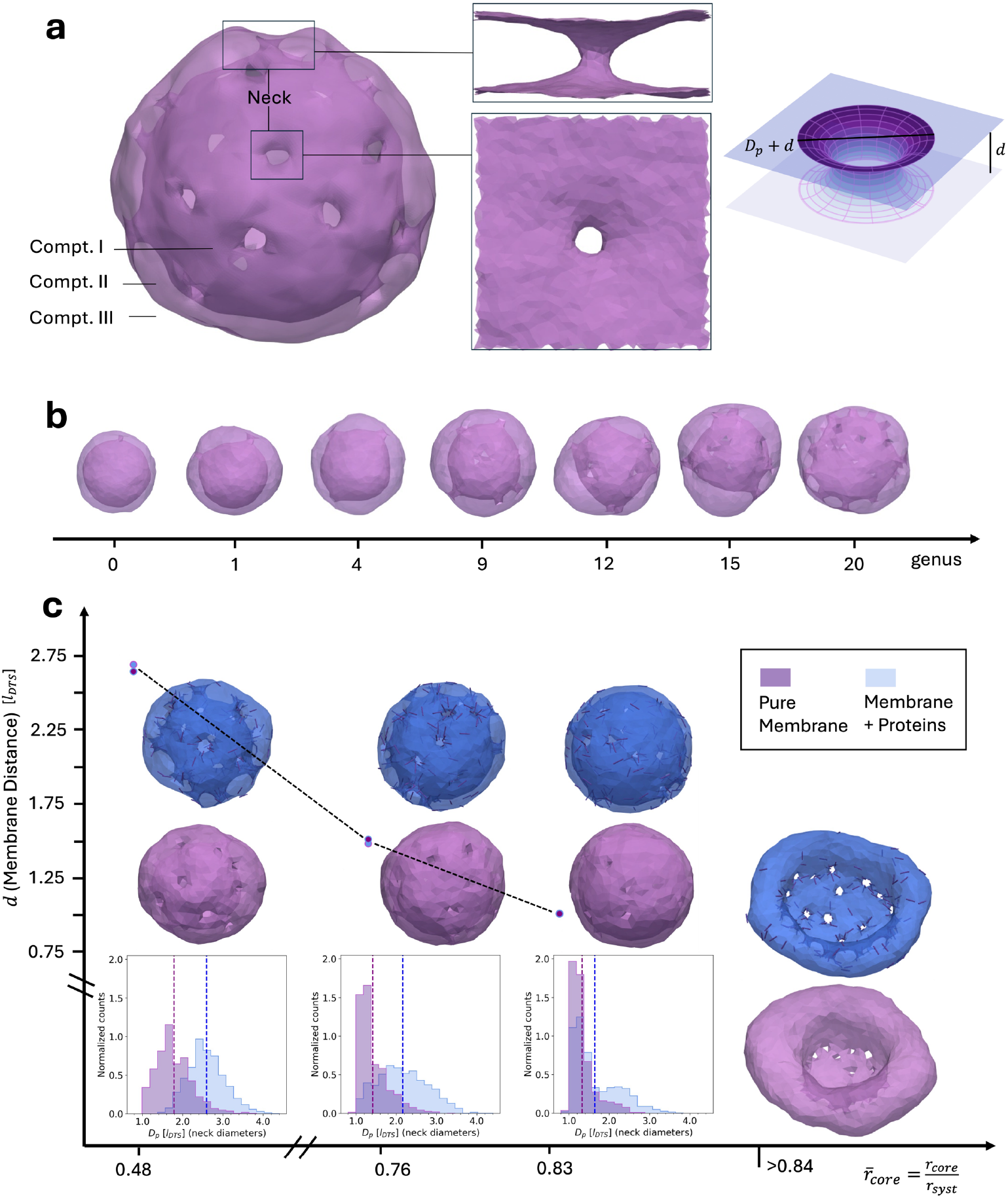
Simulations of high-genus stomatocytes. **(a)** A model of the high-genus (g=20) stomatocyte morphology (left), highlighting membrane necks and compartments created by the shape. A translucent rendering was used to make internal structures visible. Compt. I, the innermost part of the structure, corresponds to the nuclear lumen for NE, Compt. II, the volume between inner and outer membrane, represents the NE perinuclear volume. Compt. III lies beyond the outer membrane, *e*.*g*. the cytosol. The zoom-in to a membrane neck (snapshot, middle) was used to examine neck behaviour close-up in PBC in a rectangular cuboid box using DTS simulations. As a theoretical model we employed a toroidal membrane neck (right). **(b)** Stomatocyte morphologies from genus 0 to genus 20 observed after equilibration for reduced volume *v* = 0.3 at bending rigidity of *κ* = 20 *k*_*B*_*T*. **(c)** A genus-20 stomatocyte (21 necks) at *κ* = 10*k*_*B*_*T* was simulated with a continuously growing core bead to represent rising osmotic pressure in Compt. I, with and without curvature inducing proteins. For all spherical stomatocyte shapes (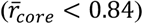), distributions of the neck diameters are shown below the corresponding simulation snapshots. They were obtained from the last 50 frames (resp. 1050 datapoints) of subsequent equilibrium simulations at constant core radius. IM-OM distance, *d*, is given as the mean of the average *d* (over the membrane) of the same 50 frames; errors are not shown as they are smaller than the datapoint markers (standard error of the mean *SE*(*d*) < 0.007*l*_*DTS*_ for all *d*).

An excellent experimental platform for exploring stomatocyte morphology in cellular context is the nuclear envelope (NE). While the NE is a highly complex system, with encapsulated DNA and intermediate filaments within the nucleus also exerting pressure on it ^24–28^, it is generally accepted that the dominant pressure within the nucleus and cytoplasm arises from the osmotic pressure of proteins and RNA molecules localised to these compartments rather than from chromatin or its associated ions ^27,29,30^. Experimental studies of the NE largely agree that increased tension on the nuclear membrane leads to an increase in the diameter of the intact nuclear pore complex (NPC), effectively dilating the nuclear pores ^31–35^. In these experiments, tension has been varied through change in osmotic pressure or energy depletion ^31,35^, by considering different states of cell differentiation ^32^ or by comparing purified NPCs with those in the native cellular context ^33,34^. At lower NE tension a constricted state has been found ^31–33^. It is commonly considered that this NPC constricted state is a ground state, from which it then dilates under increased NE tension. However, Taniguchi et al. ^32^ showed that in cells with structurally impaired NPCs, increase in NE tension led to NPC constriction relative to states of smaller tension. This unexpected opposite behaviour was explained as arising either from the inability of the nuclear membrane to properly transfer forces to defective NPCs or from tension being already alleviated by an overdilation of some NPCs^32^. To add to this contradicting behaviour, a molecular dynamics simulation of the NPC in a membrane neck also showed that NPC diameter could be maintained under tension, in addition to the expected tension-driven dilation ^34^.

Since the NPC is a complex biomolecular aggregate ^36,37^ the above diverse behaviour could arise from its intricate mechanics and mechanosensitivity. In this work, however, we provide an alternative possibility, namely that the complex mechanical response of the membrane in the associated morphology could account for the two-sided responses to membrane tension. For this, we perform mesoscopic simulations of biomembranes together with theoretical analyses to shed light on the response of membrane necks. We use the FreeDTS ^7^ software, where a membrane is represented by a dynamically triangulated surface with the basic length unit *l*_*DTS*_ being the minimal distance between any two vertices of the membrane mesh. Its shape is governed by a discretized Helfrich-Hamiltonian with additional potentials to model certain conditions, like volume constraints. Equilibrium configurations are determined using Monte Carlo sampling and details can be found in the Methods section. First, we explore a simple membrane for a more general approach and attain results transferable to any such membrane system. We then add complexity by incorporating a protein complex, addressing the discussed discrepancies and elucidating the role of the NE membrane in nuclear mechanosensing. Our findings suggest a previously overlooked material property of membranes to be of central relevance for understanding the characteristic response to mechanical stress of these membrane systems, which are at the heart of sub-cellular structure.

## RESULTS

### 1. Stomatocyte forms easily, especially for surfaces with high topological genus

We first obtained an overview of conditions in which a stomatocyte morphology forms, using volume control over a wide range of topological genera, from the well-understood spherical case (genus 0) up to a genus of 20. The systems were prepared by relaxing and equilibrating closed cuboid membrane surfaces, each with a fixed genus, *i*.*e*. a constant number of membrane necks. **Fig. 1b** shows a summary of exemplary membrane configurations for different genera for a reduced volume of *v* = 0.3 in good agreement with the previous studies ^9^. Indeed, similar results were obtained for *v* < 0.45 or a negative spontaneous curvature (results not shown). For high-genus surfaces (g = 18-20), spherical stomatocytes also formed without the need for any volume constraint or negative spontaneous curvature. This may be due to the mixing entropy contributions of the necks, as at low temperatures (or high bending rigidity), the necks concentrate around a disc that divides the inner space into two compartments (Supplementary Fig. 1b). For surfaces with smaller topological genus (fewer necks), the necks tend to concentrate in a specific region of the vesicle (see Supplementary Fig. 1a).

As a note, formation or removal of necks, *i*.*e*. changes in surface genus, generally require membrane fission and fusion processes which are subject to high energy barriers ^38–41^. Therefore, membrane topologies (here the number of necks) often remain constant while shapes vary. Here, we aim to uncover the membrane mechanics that determine membrane neck shape, most importantly size, and hence focus on systems with a fixed number of necks, *i*.*e*., constant topology.

### 2. Stomatocyte necks first constrict, then dilate under internal pressure

We then selected one of the structures with g = 20 from the previous section to investigate its response to an increase in internal pressure, which manifests as an increase in lateral tension on the membrane ^31,42,43^. In the simulation, achieving this is challenging. Unlike the well-defined volume within a triangulated surface (Compt. II in **Fig. 1a**), the internal compartment (Compt. I in **Fig. 1a**) remains connected to the environment through the necks, hence there is no unambiguous definition of its volume. Instead, we simulated the stomatocyte containing a large bead that acts as an infinite potential barrier for the vertices, meaning any shape update that places a vertex inside the bead is rejected. This effectively creates a fixed volume for the stomatocyte’s internal compartment. We do not control the lumen volume (Compt. II) at first. During the simulation, the core bead was gradually expanded, representing a slow and continuous increase in the volume of the inner compartment, analogous to the effects of internal pressure increase. The simulations were performed at near constant surface area by coupling to a harmonic stretching potential (*K*_*A*_ = 1000 k_B_T) for three different membrane bending rigidities of *κ* = 5, 10, 20*k*_*B*_*T* (see Methods for more detail and Supplementary Fig. 2 for more results). Additionally, we examined the same system decorated by 10% directional curvature-inducing proteins, which assemble in the necks; this configuration was previously used as a minimal mesoscopic model for nuclear membranes ^7^. For defined core bead sizes, we subsequently performed constant bead size simulations to obtain higher sampling and exclude the effect of the rate of bead radius increase.

In all simulations, with and without protein assemblies, the results show that increasing the core bead radius first leads to a constriction of all necks after which dilation of some necks occurs (Supplementary Fig. 2). **Fig. 1c** shows the results after additional constant bead size simulations for *κ* = 10 *k*_*B*_*T*. The constrictive response can be quantified in the neck diameter (*D*_*p*_) distribution as diameters move from larger values at rescaled core radius 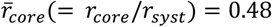 to very small values at 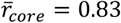. Protein assemblies in the necks stabilise larger neck diameters at small 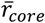, and delay and counteract constriction slightly. Only for a growing 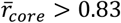 is further stretching of the envelope facilitated by dilation of most necks, both in the simple membrane and with protein assemblies (see Supplementary Fig. 2)). However, long constant core-size simulations for different core radii 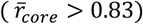 exhibit only one largely overdilated neck (see **Fig. 1c**). This indicates that multiple dilated necks are transient results of kinetics, and the lower free energy is achieved by a single overdilated configuration. If one neck overdilates all the other necks stay relatively constricted, as illustrated in **Fig. 1c** (and compared to frames from expanding core simulations, Supplementary Fig. 2).

The initial constriction in neck diameter (for 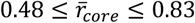) is accompanied by a reduction of over 50% in the inner-to-outer membrane (IM-OM) distance *d* in all simulations. The membrane distance starts to visibly recover and increase once overdilation occurs. These results suggest that in addition to the neck diameter the lumen volume, Compartment II in a stomatocyte (**Fig. 1a**), is another variable to be adjusted, allowing for the possibility of neck constriction by stretching. However, compared to the core volume (Compt. I) the lumen volume, due to being significantly smaller, is likely not tightly regulated by osmotic pressure (Supplementary Note 1.1). When the lumen volume is nevertheless controlled together with the core bead their effects overlap without inducing qualitatively new neck behaviour (Supplementary Fig. 3).

The constriction-dilation response also persists for membranes with spontaneous curvature (*C*_0_), representing transbilayer asymmetry effects (Supplementary Fig. 4). Notably, increased spontaneous curvature, *e*.*g*. 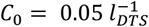, acts similarly to the proteins and slightly shifts neck diameter distributions to larger values. As the NE has a large radius (2.5 − 10 μ*m* ^44^) global membrane curvature is small (0.4 − 0.1 *μm*^−1^), and we simulated 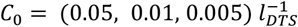, corresponding to 2.1 − 0.2 *μm*^−1^. To convert to physical units (see Methods), we used the average relaxed neck *D*_*p*_ ≈ 1.8*l*_*DTS*_ in simulations at 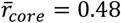 (**Fig. 1c**).

### 3. Two-state response of necks to stretching

To investigate the constriction/dilation effect further, we focused on the responses of necks in a subset of the system. Since the nuclear membrane is a large system (5 − 20 μ*m* nucleus diameter ^44^) the bending energy arising from its global spherical shape is negligible at the scale of a nuclear pore. Hence, the membrane region surrounding the nuclear pore can be approximated as locally flat. A common model used in simulations for such systems are membranes (surfaces) with periodic boundary conditions (PBC) in the XY directions, yielding results generalizable to the whole system ^45–48^. Accordingly, we consider two flat surfaces connected by a single neck with PBC, representing a small segment of the NE (see **Fig. 1a**). Such model is particularly reasonable as the density of NPCs, is relatively low in humans with 8 − 12 / μ*m*^2 49^, providing an estimated distance between NPCs of around 150 *nm* (see Supplementary Note 1.2). We present the simulation results, and afterwards consider a simple theoretical model to obtain additional intuition about these systems’ behaviours.

In PBC we first performed constant frame tension simulations ((*N, τ, T*) ensemble, see Methods and Supplementary Note 2) to account for the effect of osmotic pressure in Compt. I (see also **Fig. 1a**). Here, the frame refers to the simulation box in the XY directions (frame area *A*_*P*_ = *L*_*x*_ × *L*_*y*_), often called “projected area” in single membrane simulations and theoretical analysis ^47,48^. In this ensemble the applied tension couples the changes in the simulation box in XY direction to an energy cost. The results reveal a general behaviour: tension induced dilation only when exceeding a threshold value, while below this threshold constriction occurred. The threshold for what constituted “sufficiently high” tension depended on the bending rigidity κ of the membrane and the starting diameter of the neck. Specifically, for our starting diameter of *D*_*p*_ ≈ 8*l*_*DTS*_ in a periodic box of 30 × 30 *l*_*DTS*_^2^, we found dilation of necks for tensions (screened in steps of 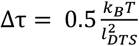 starting at 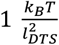 for κ = 5*k*_*B*_*T*,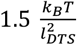 for κ = 10*k*_*B*_*T* and 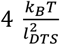 for κ = 20*k*_*B*_*T*. At lower tensions the neck underwent constriction, while at higher tensions it tended to dilate indefinitely (see Supplementary Fig. 5). When incorporating spontaneous curvature, we observed the same behaviour but lower threshold tensions (screened in steps of 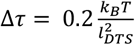 of 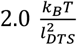 at 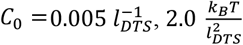 at 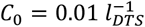 and 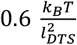 at 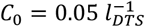 for κ = 20*k*_*B*_*T*.

To further characterise the response, we performed constant frame area ensemble simulations (*N, A*_*p*_, *T*) for a wide range of frame areas (for details see Methods). Because (positive) tension favours increasing *A*_*p*_ this approach allows us to track its effect on neck size in a more controlled manner, while sampling configurations that might be inaccessible in constant-tension simulations, where the area can expand too rapidly. The resulting average bending energy as a function of *A*_*p*_ is shown in **Fig. 2a** for three different bending rigidities 5, 10 and 20 *k*_*B*_*T*. For all the depicted cases, we first observe a decrease in the mean bending energy with increasing frame area. Then, upon reaching a branching *A*_*p*_, some replicas remain in the low-energy minimum, while others settle into a local minimum with higher energy. This marks the transition point or region, highlighted in the inset of **Fig. 2a**: the branching is preceded by a point of degeneracy, where replicas’ equilibrium energies cover a range of values. Then, replicas equilibrate at either very low or higher energy states. Beyond a critical *A*_*p*_, configurations in the lowest energy minimum disappear and the energy increases approximately linearly with frame area. The frame area at which the branching takes place is higher for larger bending rigidity. Conversely, a higher area compressibility (see Method section) pushes the branching point to smaller frame areas (**Fig. 2b** and Supplementary Fig. 7).

**Fig. 2:**
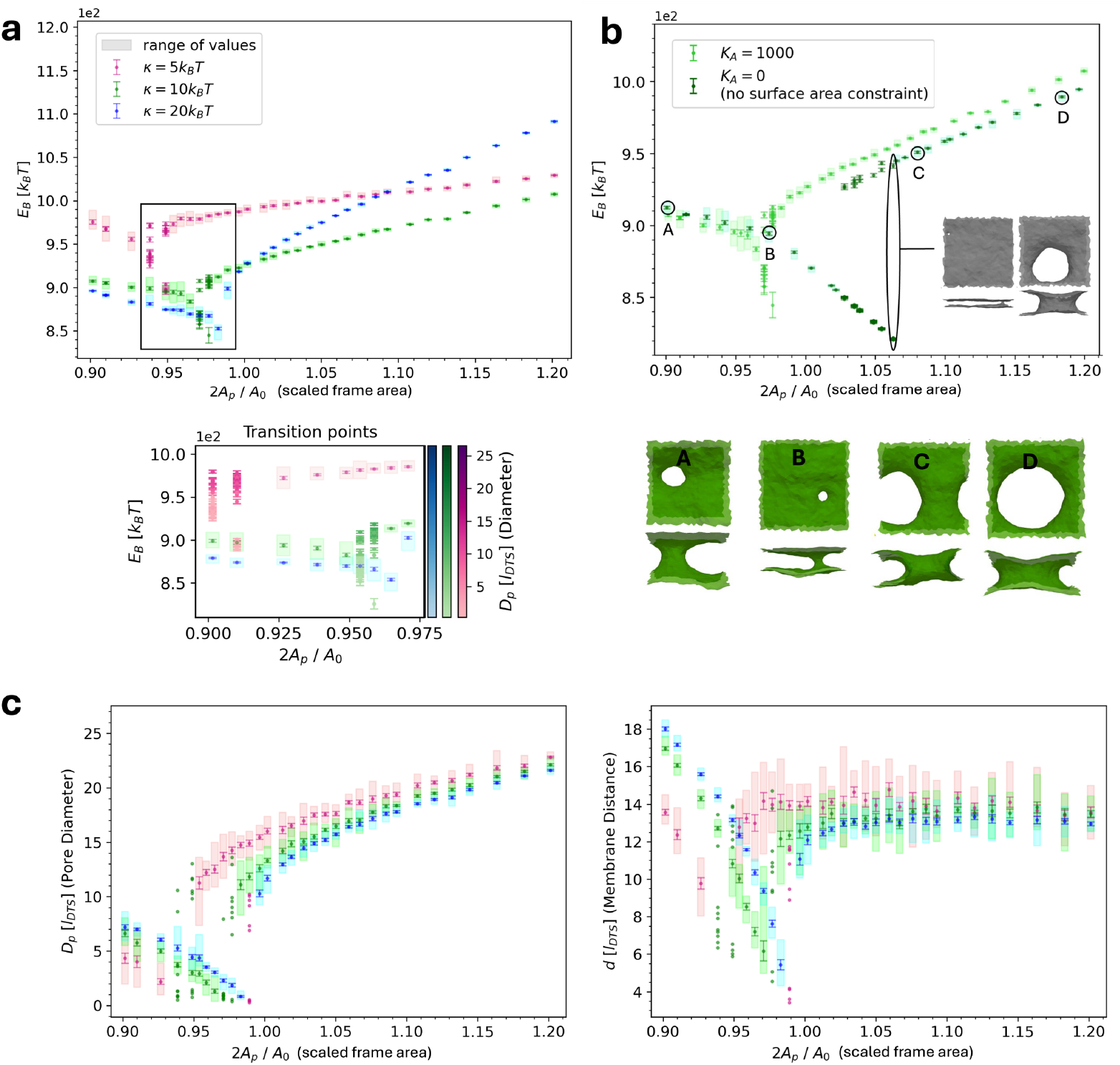
Constant frame area simulations of double membranes connected by a neck in PBC. **(a)** Bending energy as a function of frame areas for different bending rigidities. Inset created from additional 30 replicas showing their individual behaviour at transition points, where the energies are related to observed diameters *via* a colour scale. Results were obtained from simulations with near-constant total surface area (*K*_*A*_ = 1000 *k*_*B*_*T*). For each parameters set and at each frame area 10 replicas were run (30 replica at transition points for inset). Each datapoint corresponds to their mean energy, errorbars indicate the standard error of the mean. At transition points in frame area some replicas show high, others low bending energy, indicating multiple local minima. Hence, all replicas are displayed instead of the mean; an error on each replica was obtained by block averaging over the last 10^6^ MC steps after equilibration of each simulation (20 blocks, length of 500 energy outputs; energy was saved every 100 MC steps). **(b)** Influence of area compressibility *K*_*A*_ (see also Supplementary Fig. 7) and simulation snapshots of final morphologies. Results are shown for *κ* = 10 *k*_*B*_*T* at near-constant surface area (see (a)) vs. no surface area constraint. Snapshots are shown for *K*_*A*_ = 0 *k*_*B*_*T* but are representative for general behaviour, *i*.*e*. for all area compressibilities. **(c)** Neck diameters (left) and membrane distance (right) after equilibration of one membrane neck in PBC at different frame areas. Results were obtained from simulations with near-constant surface area (*K*_*A*_ = 1000 *k*_*B*_*T*). Datapoints correspond to mean values of 10 replicas and errors were obtained as the standard error of the mean. Individual replicas at transition points are shown without errors. Note: while vertex distances always exceed 1 *l*_*DTS*_, the analysis method may yield smaller values for extremely narrow necks. The procedure of obtaining neck sizes is detailed in the Methods section.

As shown in **Fig. 2a-d**, characteristic changes in neck size go along with the described energetic behaviour. While early increases in frame area are facilitated by neck constriction and decrease of IM-OM distance, greater increases of *A*_*p*_ lead to neck dilation and relaxation of the IM-OM distance. The decrease of bending energy at small frame areas can be attributed to neck constriction, but also to smoothing of the membrane far from the neck site (see snapshots **Fig. 2b**). At branching points, neck diameter dilation, corresponding to the higher-energy local minimum, as well as constriction, corresponding to the lower-energy minimum can occur. It should be noted that the coexistence of the two minima despite one having significantly higher energy, could indicate an entropic preference for the dilated system compared to the highly constricted system. This is supported by the observation that the membrane becomes increasingly flat in the constricted system, suppressing many undulation modes (see **Fig. 2b**).

Overall, the neck diameter and IM-OM constriction followed by (over)dilation agrees with our observations in the stomatocyte simulations (see above). It becomes apparent that one key component to mechanistically explain the two-phase behaviour is the counteracting interplay of bending energy with the demand for larger frame area. We note that at effectively constant surface area *A*_0_ (*K*_*A*_ = 1000 k_B_T), the transition always takes place at 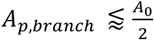. This indicates that constriction occurs as long as the membrane surface area is large enough to span the specific frame area. Therefore, it can also explain why the transition point is shifted by bending rigidity and area compressibility. Higher bending rigidity increases available membrane surface area by smoothing undulation modes, while higher area compressibility resists surface extension. Hence, larger or smaller *A*_*p*,*branch*_ can be facilitated, before the system reaches the physical constraints of how far it can stretch without neck dilation.

### 4. A simplified theoretical model predicts and characterizes constriction & dilation

To investigate the underlying effects further, we developed a theoretical model. Consider a region of the membrane around a neck in which beyond a certain cut-off radius 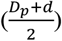 from the neck centre the membrane becomes flat (**Fig. 1c**). Thereby the neck is embedded in two square pieces of membrane with side length *L*, corresponding to IM and OM (inner and outer nuclear membrane in case of the NE). We assume that the neck is toroidal with diameters *D*_*p*_ and *d*, defining the smallest part of the channel and half of the IM-OM distance, respectively. For an evaluation of this assumption see Supplementary Note 5 (and Supplementary Fig. 19, 20). The bending energy of the system is described by the Helfrich Hamiltonian,

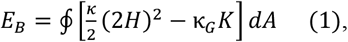

with bending modulus κ, and gaussian modulus κ_*G*_. The Gaussian term (second term) is constant if there is no topological change to the surface and therefore will be neglected. Then the bending energy of the toroidal neck is

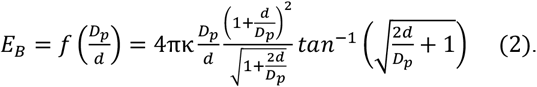

A lateral tension is represented by an energetic contribution *E*_τ_ = −*τA*_*p*_ with tension parameter *τ* and frame area *A*_*p*_. Under the constraint of constant total surface area *A*_0_, the rescaled energy (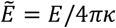) will only depend on 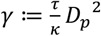 and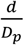.We minimised this energy numerically (details in Supplementary Note 3). For zero frame tension (*γ* = 0) the bending energy has a minimum at 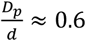 in agreement with other studies on toroidal pores ^50,51^ (Supplementary Note 3, Supplementary Fig. 8a). This points to a degeneracy of the neck size for tensionless membranes which qualitatively matches with our simulation results (Supplementary Fig. 8b). Non-vanishing frame tension, however, leads to degeneracy lifting. There,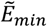 possesses a single local maximum, yielding an unstable critical point γ_crit_ ≈ 10.667. This means that for systems with *γ* < γ_crit_ (*γ* > γ_crit_), at a given tension and bending rigidity, a membrane neck constricts (dilates) fully due to the tension (**Fig. 3a**). In other words, for a given neck diameter there exists a critical tension: below that value, the neck constricts, and above it, the neck dilates. This result is in good qualitative agreement with our simulations results. Moreover, γ_crit_ = *τ*_*crit*_ *D*_*p*_^2^/*κ* ≈ 10.667 provides a very simple but strong equation to assess the system behaviour that also agrees well with critical tensions recovered from simulations (see Supplementary Fig. 5). Assuming the purified NPC diameter as our neck diameter *D*_*p*_ = 43 *nm* ^33,52^ and for realistic membrane bending rigidities in the range of 3 − 15 × 10^−20/^*J* ≈ 7 − 35*k*_*B*_*T* ^53^, our model predicts critical tension parameters of 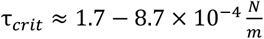 NE tension has been estimated at about 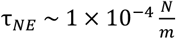, an order of magnitude higher than typical plasma membrane ^54–56^. Thus, the critical tension lies within the physiological range of the NE tension, making the described constriction-dilation phenomenon highly relevant for NPC responses to mechanical stimuli.

**Fig. 3:**
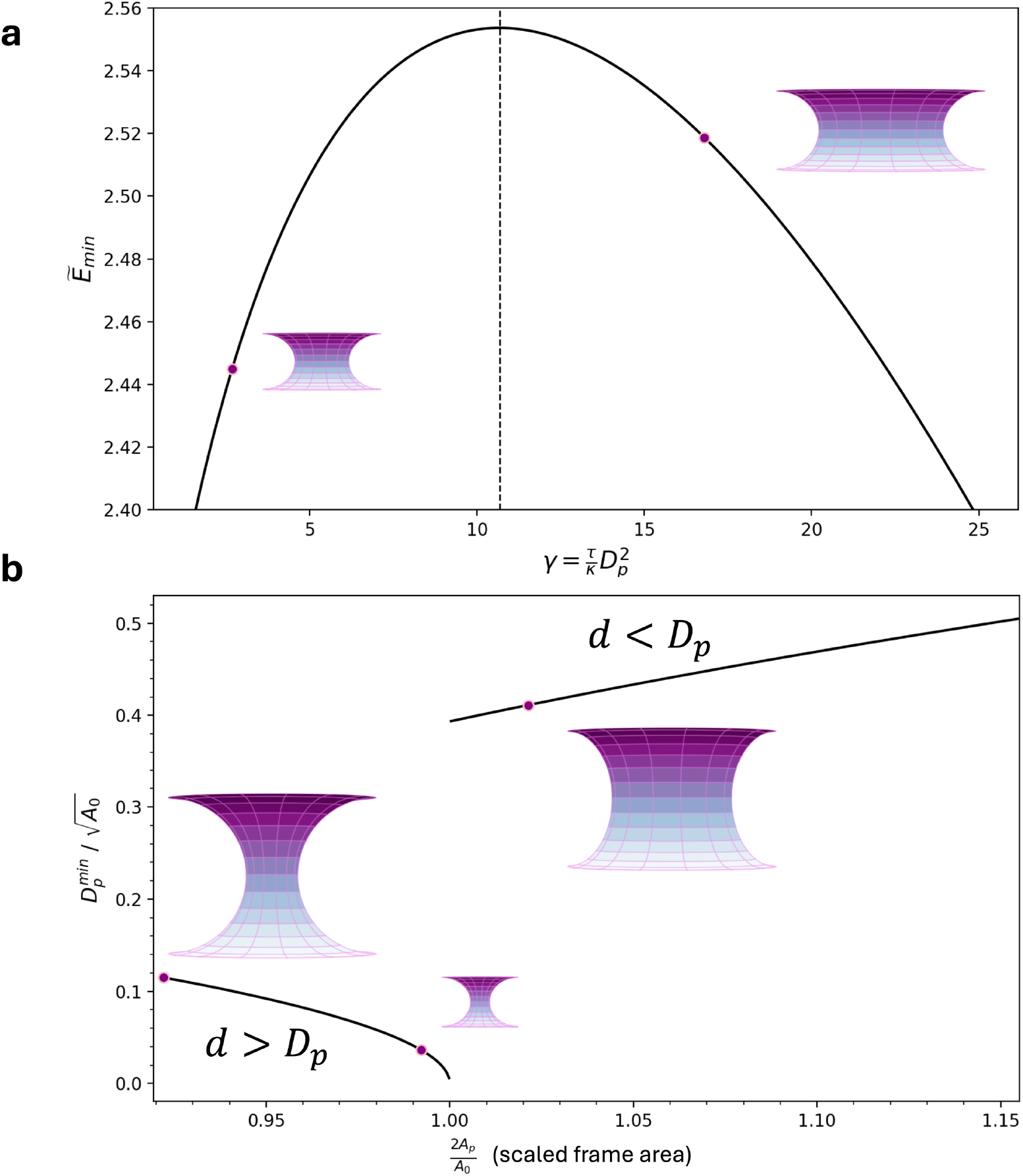
Theoretical model predictions. **(a)** Minimum rescaled energy 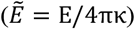 for a toroidal neck under tension and under constraint of constant surface area, as a function of dimensionless parameter *γ*. An energy barrier divides regions where constriction or dilation would be favoured at a specific tension and bending rigidity. A schematic view of the neck is shown. **(b)** Neck diameters *D*_*P*_ corresponding to minimal bending energies (see also Supplementary Fig. 11) under constraint of constant frame area over a range of frame area values. Additionally, surface area was controlled *via* a harmonic potential with target surface area *A*_0_ (*K*_*A*_ > 0). The neck diameters are scaled with the target area and are independent of bending rigidity. A regime of constriction, characterised by neck diameters smaller than the membrane distance *d*, is followed by dilation, where neck diameters are larger than the membrane distance. In the dilation regime, the neck diameters have finite sizes, as they are geometrically constrained by the system size, but they represent infinite dilation. This is because no local energy minimum exists once 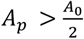, *i*.*e*. when neck diameters are larger than the membrane distance (*D*_*P*_ > *d*).

The toroidal neck model is based on simplifying assumptions that facilitate understanding but has limited applicability when additional complexity is introduced, such as spontaneous curvature *C*_0_ or pressure difference (*P*) between the lumen and outer environment. These effects could lead to configurations far from the model’s assumptions and may therefore only be valid in the small-perturbation regime. Hence, one should rely mostly on the simulation results for these more complex cases, but the analytical extensions can be found in Supplementary Note 3. For a small spontaneous curvature, the bending energy *E*_*B*_ changes accordingly (results Supplementary Fig. 9). A small pressure difference due to the lumen volume (*V*) can add a term −P*V* to the energy (results Supplementary Fig.10). Both modifications cause only minor changes to the predicted constriction-dilation behaviour. Notably, *C*_0_ > 0 shifts the threshold position γ_crit_ downward in agreement with the lower threshold tensions in simulations (results 3.).

To evaluate the relationship between simplified theory and simulation results further, we solved the theoretical model for constant frame area (derivation in Supplementary Note 4). **Fig. 3b** shows the theoretical prediction of the neck diameters over a range of frame areas, similar to the simulation results in **Fig. 2c**. Unlike the simulations, the theory predicts that the neck diameter is independent of membrane bending rigidity (the minimised energy is simply proportional to κ). However, we see very good agreement in the general behaviour of the predicted diameters with those obtained from simulations: while an increase in *A*_*p*_ first leads to constriction with increasingly negative slope, a transition point occurs, at which there is a jump to dilated states. From there on the diameter increases with frame area. Before the transition point is reached, an increase in frame area is facilitated by increasing constriction of neck diameters and IM-OM distance. This process is governed by bending energy in our model. It predicts a constant minimal bending energy value at all frame areas in the constrictive region, which depends on the bending rigidity (Supplementary Fig. 11c). The quantitative deviation of theory results (**Fig. 3b**) from the simulations (**Fig. 2a**) likely arises from the influence of thermally induced membrane undulations in the simulations. The transition occurs at 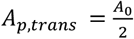. Above this value, 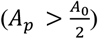, there is an asymptotic convergence of the minimizing diameter ratio from above to 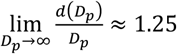, while the bending energy as a function of both diameters exclusively minimises at 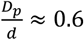. Therefore, after the transition 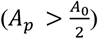 the energy decreases with increase in *D*_*P*_ until the membrane size limit is reached. The diameters are hence not energetically constrained, but instead determined by the geometry of our setup, so 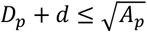. This limit leads to an approximately linear increase in *D*_*P*_ very similar to the one seen in simulations with large frame areas. The theoretical limit is not suitable to estimate the diameter values observed in simulations accurately (compare **Fig. 2a**). Likely, the difference is again caused by membrane undulations in the simulation, which lead to stricter constraints on *D*_*P*_. Fitting the approximately linear energy increase after the transition point in simulations and comparing to the theoretical prediction shows that the ratio of the slopes for different bending rigidity agree well (for details see Supplementary Note 6), implying that the simulations in the dilation regime are predominantly steered by the same physics captured in the theory.

### 5. Multiple Neck Systems also show two state behaviour

Next, we examined how a system containing multiple necks in PBC responded to increasing frame area. For this purpose we chose systems of two, four and six necks per membrane in PBC. **Fig. 4a,b** (and Supplementary Fig. 12) shows the average bending energies after equilibration as a function of frame area *A*_*P*_ for different bending rigidities of 5, 10 and 20 *k*_*B*_*T*. (Here we have used area compressibility *K*_*A*_ = 1000 *k*_*B*_*T*). The multiple neck systems energetically behave very similar to the single neck system. For small increases in frame area, the bending energy decreases until a transition point is reached, after which it increases. The frame area at which the transition takes place increases with increasing bending rigidity. By inspecting the simulation snapshots, we find that this is also caused by constriction of all necks during small growth in frame area. For larger frame areas beyond the branching point, the system exhibits only one dilated neck. If additional dilated necks are present in the initial configurations (taken from simulations under tension), they transition to a more relaxed, constricted state during constant *A*_*P*_ simulation. This suggests that two or more dilated necks are transient features resulting from fast stretching dynamics, rather than representing the system’s equilibrium configuration. The overall similarity to single-neck systems confirms that the underlying principles governing neck size in response to increasing frame area remain consistent across systems with different numbers of necks.

In systems with 6 necks, we did nevertheless see indications of multiple transition points occurring. These systems again showed constriction of all necks before the first transition point, and dilation of only one neck after it. However, this was followed by later dilation of more necks at larger frame areas (see **Fig. 4a,b)**. At the second possible transition point in the data for κ = 10*k*_*B*_*T*, the snapshots from the simulations highlight that the replicas with fewer dilated and more constricting necks have lower bending energy. This helps to explain the observation that only one dilated neck was found in equilibrated systems of genus two and four, and earlier in stomatocyte vesicles (see also **Fig. 1c**). Taken together, it seems to be more energetically efficient to have one neck dilated by a larger amount than two (or multiple) necks dilated by a smaller amount each. The energetic cost of dilation of more than one neck at larger frame areas seen in the 6-neck system may be compensated by an entropic gain. While multiple-neck systems behave similarly to the single-neck system overall, their response is more complex. Significant degeneracy may exist within the dilation regimes; further exploration of these feature is beyond the scope of this work.

**Fig. 4:**
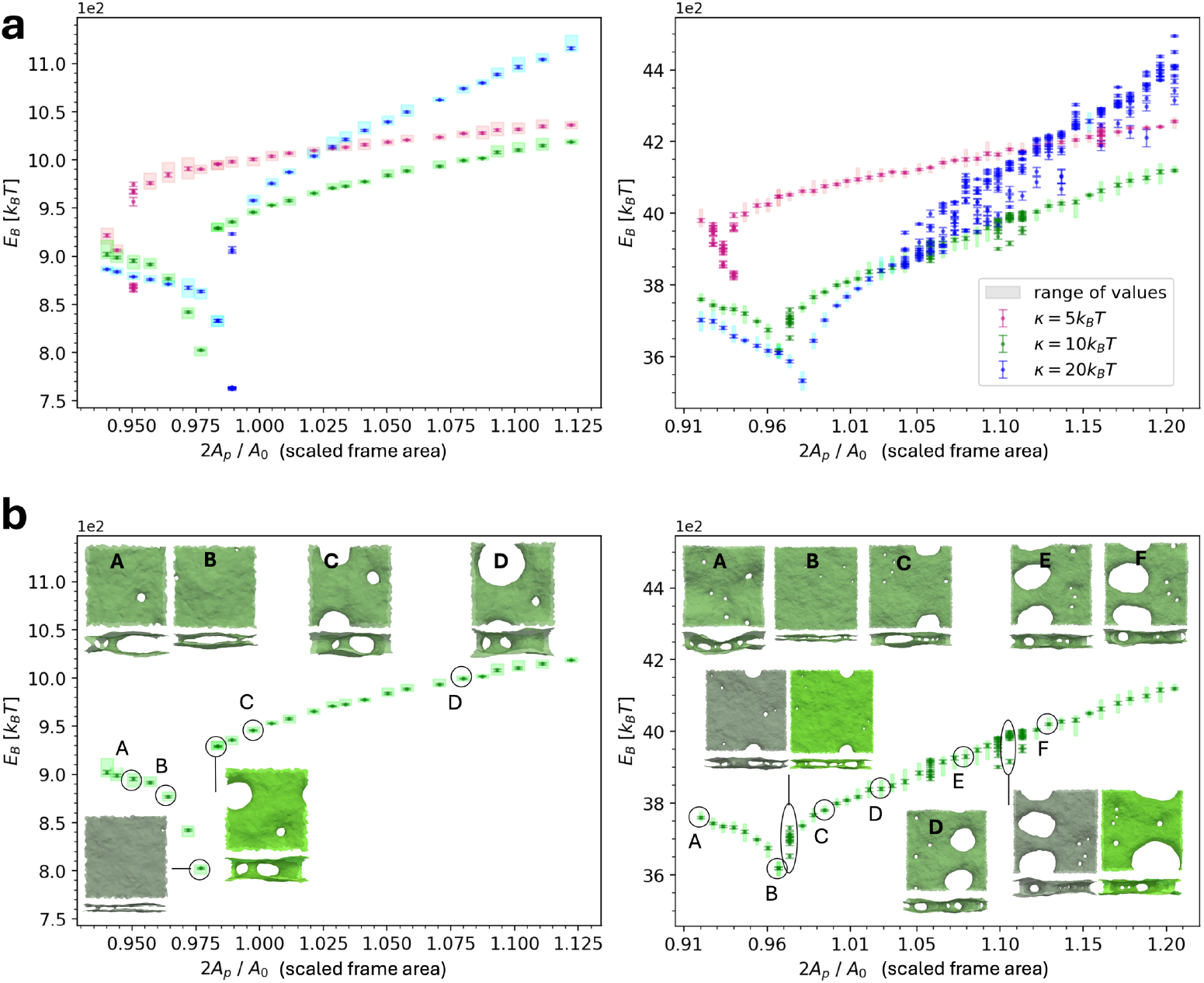
Systems with multiple necks in PBC. **(a)** Bending energy after equilibration of two (left) and six (right) membrane necks in PBC at different frame areas. Results were obtained from simulation with near-constant surface area 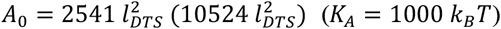 (*K*_*A*_ = 1000 *k*_*B*_*T*). While two necks (and four necks, see SI) show one transition point like a single neck, at six necks multiple such transitions exist, possibly overlayed. The occurrence depends on the bending rigidity. At each frame area 10 replicas of the simulations were run. Each datapoint corresponds to their mean energy and errorbars indicate the standard error of the mean. At (possible) transition points in frame area, all replicas are displayed instead of the mean; an error on each replica was obtained by block averaging: for two (six) necks over the last 10^6^MC steps (8 × 10^5^ MC steps) of each simulation after equilibration (20 blocks (16 blocks), length of 500 energy outputs; energy was saved every 100 MC steps). **(b)** Visualisation of the necks’ behaviour for two (left) and six (right) necks in PBC at bending rigidity of *κ* = 10 *k*_*B*_*T*. This illustrates a second transition point or region in the six-neck-system, facilitated by successive dilation of more than one neck.

### 6. Protein Addition to the Neck Retains the Key behaviours

Finally, we tested systems containing a neck decorated by a protein complex (for details, see Methods). We employed a minimal model for the protein complex, using elongated inclusions with a preferred negative curvature that assemble into the neck, effectively forming a protein complex (see **Fig. 5**). By varying attractive protein-protein interactions, we explored different stability of the complex structure. This model implicitly mimics the complex’s interaction with the membrane. The system was initially equilibrated in a tensionless state of the membrane, to allow for proteins to assemble in the membrane neck. Then it was subjected to a range of frame tensions under the constraint of near-constant total surface area (*K*_*A*_ = 1000*k*_*B*_*T*).

**Fig. 5:**
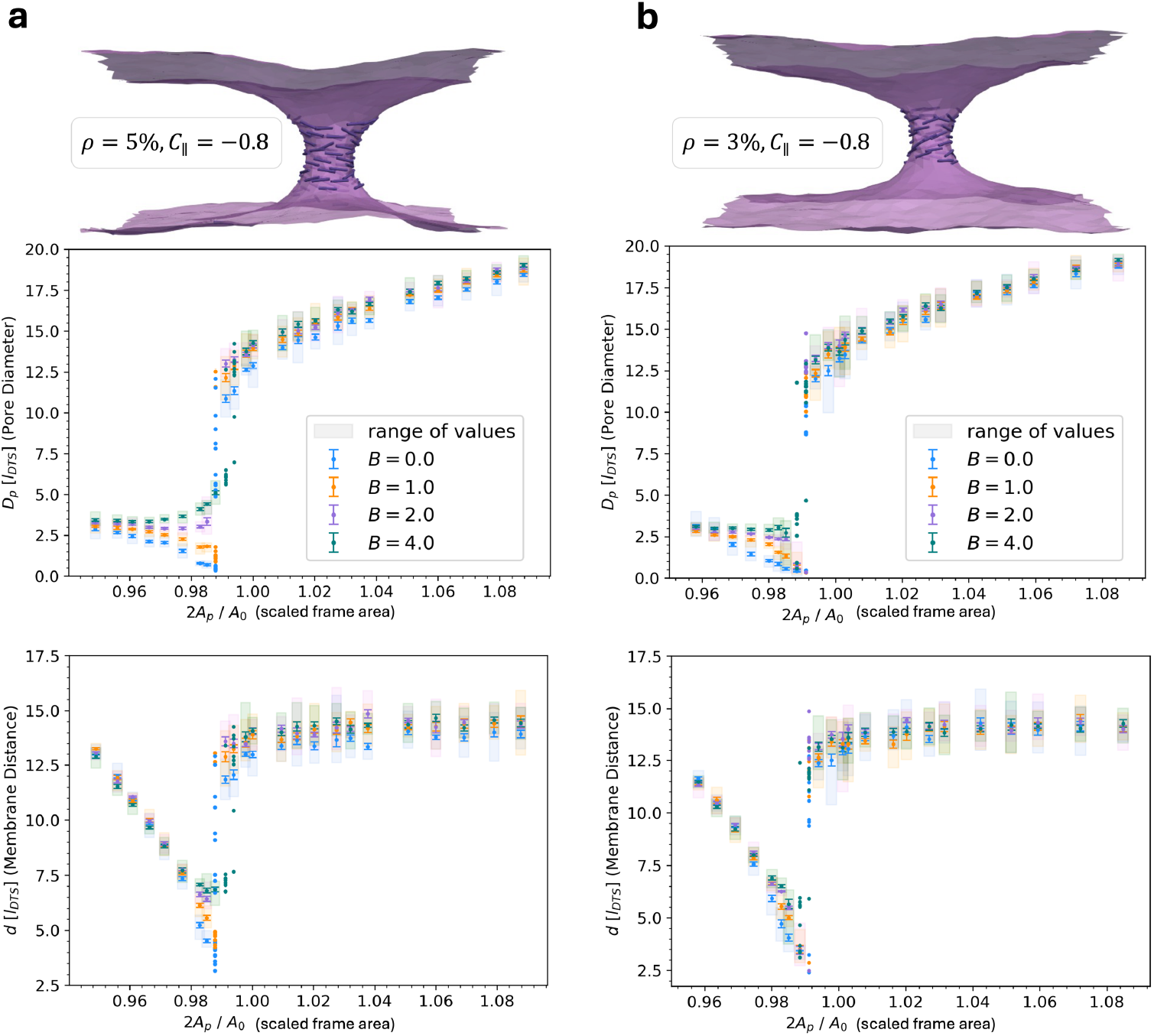
Constant frame area simulations of double membranes connected by a neck with protein assembly in PBC. Neck diameters (top) and membrane distances (bottom) of simulations over a range of constant frame areas, for different protein complex sizes of **(a)** a coverage of *ρ* = 5% of membrane vertices with an inclusion, and **(b)** a coverage of *ρ* = 3% of membrane vertices with an inclusion, *i*.*e*. a protein. The protein interaction strengths were varied (*A* = 1*k*_*B*_*T, B* in *k*_*B*_*T* as specified in legend) at constant curvature preference (*C*_∥_in 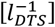). Results were obtained from simulations with near-constant surface area (*K*_*A*_ = 1000 *k*_*B*_*T*). Datapoints correspond to mean values of 10 replicas and errors were obtained as the standard error of the mean. Individual replicas at transition points are shown without errors. The procedure of obtaining neck sizes is detailed in the Methods section. The proteins are depicted as lines to indicate their orientation on the membrane.

We first examined how protein abundance, *i*.*e*. size of the complex, induced curvature, and interaction strength influence the response of membrane neck size to increasing tension (see also Supplementary Fig. 13). A larger complex with strong protein interactions suppresses neck constriction at lower tensions and causes a controlled slight neck dilation at increased tension. Compared to smaller protein complexes with less interaction, it surprisingly causes earlier occurrence of overdilation for 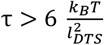. Increasing the induced curvature (from 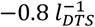 to 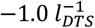) or decreasing the interactions between proteins prolongs neck stability (overdilation at higher tension). A lowered interaction strength furthermore allows for a notable constriction response to increasing tension. Decreasing the size of the protein complex as well leads to an even stronger constriction, accompanied by neck stability for at least 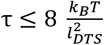. Taken together, this shows that the proteins retain the key membrane responses to tension but counteract and control them increasingly the larger and/or stiffer the protein complex is.

We studied the response in detail by performing constant frame area ensemble simulations, and examining the behaviour of energy (Supplementary Fig. 14) and neck diameters (**Fig. 5**) as a function of frame area for the large protein complex (5% coverage), as well as for a smaller protein complex (3% coverage) with varying interactions. This underlines the observations under tension, by showing that the large, cohesive complex can indeed suppress constriction, especially through strong interaction between the proteins. Compared to a simple membrane, the complex first stabilises a moderately small neck, while the IM-OM distance decreases (see **Fig. 5**), until dilation sets in at larger frame areas. This dilation can occur more continuously compared to the abrupt behaviour seen in a pure membrane. A less cohesive or smaller protein complex instead retains all key behaviours of the pure membrane, displaying constrictions for smaller frame areas followed by dilation after 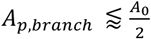. The constriction and dilation are more pronounced and the transition more abrupt the weaker the interactions between proteins are. Neither the protein abundance nor the interaction strength largely influence the location of the transition point (Δ(2*A*_*P*,*branch*_ /*A*_0_) ⪅ 0.008 for all comparisons).

## DISCUSSION & CONCLUSION

Stomatocyte shapes are highly relevant in both biological and synthetic membranes. They can divide space into three distinct compartments. Two of these compartments are connected *via* narrow necks, which can serve as intriguing sites for the assembly of specific biomolecular complexes to regulate material transport between them. Understanding how the size of these necks responds to external stimuli is therefore important to elucidate cellular processes (*e*.*g*. nuclear mechanosensing). In this work, we used mesoscale computer simulations and theoretical analysis to investigate this behaviour. Our findings indicate that at low membrane tension or small pressure differences between the innermost and outer compartments, the necks constrict. Conversely, at higher tension, they dilate and open. Parameters such as bending rigidity and area compressibility govern the transition between these two phases. The two-phase behaviour arises solely from the mechanical properties of the membrane in the stomatocyte configuration, particularly in surfaces with high topological genus. It can be modified by the presence of protein assemblies, stabilising against constriction and dilation to some degree depending on their properties (induced curvature, interaction strength, size of the assembly), but the key two-phase tension response remains. In addition to this non-intuitive mechanical response, necks in membranes with high topological genus exhibit highly mobile positions ^8,57,58^, especially for necks with smaller diameters (see Supplementary Fig. 15, Supplementary Video 1).

As observed in our constant frame area simulations, the frame size, which relates to lateral tension, can control the transition from narrow to wide neck states. Such bimodal response to mechanical stimuli can be a more general phenomenon in membranes. For example, in a geometric setting orthogonal to ours, vesicles connected to membrane segments also exhibit two-state transitions: analogous to tether pulling, applying the stimulus along the connecting neck’s axis to vary its length caused a shape switch from catenoidal necks to narrow cylindrical tubules ^59^.

The two-state response to lateral tension may explain several experimentally observed behaviours, especially the dilation of intact NPCs but also the constriction of defect NPCs under tension. Our results suggest that larger membrane necks or nuclear pores require a lower threshold tension for them to dilate, making them less restrictive on traffic through them. Since our model predicts threshold tensions in the physiological range, constriction too should play a role in the nuclear pores’ adaptation to mechanical stress. It could then *e*.*g*. explain the unexpected observations in cases of defect NPCs ^32^, challenging the assumption that dilation is the sole response to internal (osmotic) pressure or lateral tension. While the influence of the protein complex was not explicitly modelled, our results support the notion that an intact NPC safeguards nuclear pores against overdilation ^32^. We also propose that the nuclear membrane itself transmits the forces to the NPC, as, according to our findings, the behaviour of the simple membrane alone supports the previously reported NPC adjustments to changes in tension. Therefore, other potential sources of force transmission, like the LINC complex ^60^, might not be needed. However, the NE is far more complex than our minimal model, which is designed to capture only the generic behaviour. Firstly, the NPC is an assembly of proteins that interact in a complicated manner which is not fully captured in our model. Secondly, the NE is covered by several groups of proteins, which additionally interact with nuclear lamina and chromatin ^61–63^, adding complexity to the membrane mechanics. Hence, we expect our results here to represent only the qualitative and fundamental behaviours of the NE and NPC system. Indeed, for a more detailed understanding, methods such as multiscale simulations ^64,65^ that incorporate the details of the molecular systems while reaching the relevant time and length scales are needed. This might also include a more detailed comparison to experimental results and combining simulation and experimental studies to complement each other ^6^. Yet, the generality of our model is also a key strength. While it does not capture fine-grained details, it demonstrates robustness against variations in the chosen force field and potential unaccounted effects, suggesting the universality of the observed behaviour.

Neck dilation in stomatocytes is fundamentally different from that of spherical closed vesicles. It is governed by distinct physical mechanisms and results in different features. In spherical closed vesicles, strong pressure differences cause the membrane to stretch. Once a critical tension is exceeded, a pore forms, which can eventually develop into a hole if the pressure release is too slow. The stretched membrane then relaxes into a tensionless state, allowing the hole to expand further and ultimately compromise the vesicle’s structural integrity. In contrast, neck dilation in stomatocytes occurs without pore or hole formation, and the membrane remains intact. Moreover, the tension required for neck dilation in stomatocytes is significantly lower (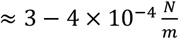 for κ ≈ 15*k*_*B*_*T*, see above) than that needed for hole formation in vesicles (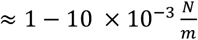, membrane rupture tension ^66,67^). This indicates that stomatocytes offer a finely tuned mechanism for compartmentalisation, providing a level of precision different from the commonly used amphiphilic assemblies like bilayers. This distinctive feature highlights the potential of high-genus membranes as functional units in synthetic biology and membrane engineering.

Since our results are based on a very generalised system, the findings could extend to other membrane systems with necks under tension, like the endoplasmic reticulum (ER). Interestingly, simulations with neck (over)dilation in PBC gave stretched configurations that had formed three-way junctions characteristic of the ER structure (see Supplementary Fig. 16). ER three-way junctions are argued to arise from a substantial membrane tension ^68,69^, making it unsurprising that our observations may be applicable to such systems. In turn, computational studies on ER membranes during transformation of junctional knots to regular junctions have shown an interesting similarity to the neck behaviour in stomatocytes. There, small membrane necks (also referred to as pores) constrict in the centre of an ER junctional knot under tension ^70^. This points to the relevance of the mechanism of membrane neck dilation and constriction in other cellular structures and processes apart from mechanosensing of the nuclear envelope.

Lastly, bridging back from ER to NE, our results may offer further understanding to postmitotic NPC assembly, which was previously reported as a sequence of membrane neck constriction and dilation events. Initially, fenestrated ER sheets’ holes (equal to largely overdilated membrane necks) shrink. Then NPC precursors accumulate in these small necks and dilate them during complex assembly and maturation ^71^. Based on our results, it stands to reason that membrane lateral tension could play a significant role in this process. Initial ER neck constriction may be facilitated by membrane tension alongside ER proteins actively driving the reshaping. Our findings on protein assemblies in necks furthermore support the idea that the precursor NPC accumulation stabilises the ER–NE hole shrinkage ^71^, allowing for prepores (preNPCs) to dilate to final size as they mature: As one can assume that prepores correspond to small protein assemblies with weaker interactions, these would counteract tension-driven constriction only slightly. Maturing NPCs then become larger and more cohesive. That could counteract constriction and possibly drive the neck diameter beyond the threshold, after which membrane tension would cause dilation. This is ultimately stabilised by the fully assembled NPCs when the final size is reached.

In conclusion, our study reveals a previously overlooked, general material property of high genus stomatocyte membranes. We demonstrate that internal osmotic pressure (or lateral tension) can lead to membrane neck constriction at low and dilation at high tensions. These results have implications not only for the nuclear envelope but also for other biological systems with (many) membrane necks. This includes the high-genus membranes of other cell organelles, like the ER, mitochondria and Golgi apparatus.

## METHODS

The mesoscopic simulations were performed using Dynamically Triangulated Surface simulations that have been shown to be very effective for simulating cell membranes at large scales ^72–74^. There are a few different models and software packages available for performing mesoscale simulations ^75–79^. Here, we have used FreeDTS ^7^, which has shown to reproduce correct thermodynamical behaviours of lipid bilayers such as the undulation spectrum ^45,80–82^. FreeDTS represents the membrane as a surface of *N*_*v*_ vertices, *N*_*e*_ edges and *N*_*T*_ triangles. The software uses the shape operator formulation to calculate membrane curvature at each vertex ^45^. A vertex is assigned with a unit normal 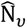, surface area *A*_*v*_, principal curvatures (*c*_1*v*_, *c*_2*v*_) and principal directions (X_1_(*v*), X_2_(*v*)) *via* a set of discretised geometrical operations. The bending energy of the surface (*E*_*B*_) is given by a discretised form of the Helfrich Hamiltonian in terms of the mean 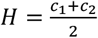, and Gaussian curvature, *K* = *c*_1_*c*_2_ as:

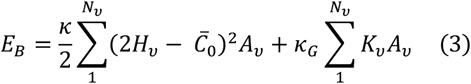

with the bending modulus *κ*, Gaussian modulus *κ*_*G*_, and spontaneous curvature 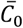. The latter represents a possible asymmetry between the two monolayers (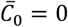 for a symmetric membrane). The simulations are evolved using a metropolis algorithm. A Monte Carlo (MC) step corresponds to *N*_*v*_ attempts to move a vertex and *N*_*e*_ attempts to flip links.

Constant tension simulations in PBC employ a position rescaling algorithm to adjust the box size ^45^. For every X^th^ Monte Carlo (MC) step, an additional box change attempt is made. This algorithm couples the system energy to

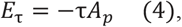

where *A*_*P*_ is the frame area.

An area constraint algorithm can be added to control the surface area of a membrane, since due to flexibility of the edge length, the area of each triangle can vary and the membrane surface is stretchable. In that case, the energy is coupled to an additional term of

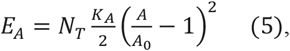

with the targeted surface area *A*_0_ and the compressibility modulus *K*_*A*_ in units of energy, such that the energy term scales extensively with the system size. For constant volume simulations, the system energy was coupled to a potential of:

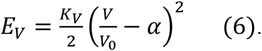

The unit length in FreeDTS, *l*_*DTS*_, can be converted to physical units *via* a known feature of the biological system under investigation. In the present work, we convert it based on the average diameter of a nuclear pore, *D*_*P*_ = 43*nm* ^33,52^. To do so we equate it with the neck diameter recovered from simulations of the tensionless system, which in PBC is D_p,sim_ ≈ 8*l*_*DTS*_ (see also Supplementary Note 5) and in the full genus-20-stomatocyte is D_p,sim_ ≈ 1.8*l*_*DTS*_ (see results 2.). This yields *l*_*DTS*_ ≈ 5.4*nm* and *l*_*DTS*_ ≈ 23.9 *nm* in each system respectively.

### Constant Frame Area versus Constant Frame Tension Ensembles

Simulations of a segment of the whole NE in PBC have been performed in two different ensembles:

#### 1. Constant Frame Tension Ensemble (*N*, τ, *T*)

In this ensemble an external mechanical tension is explicitly applied through the energy term − τ*A*_*P*_ (eq. 4). The frame area (projected area) of the membrane, *A*_*P*_, is dynamic and adapts through a specific MC move ^45^. Once the simulation is performed and the system reaches a minimal free energy, the average internal stresses from the internal energetic terms, e.g. from stretching and bending, balance those externally imposed by mechanical tension τ. This means *A*_*P*_ fluctuates around a balance of stresses (which become equal on average). A similar method has been used for simulations of particle-based membranes ^83^. This ensemble allows to probe the response of membrane necks to a specific amount of externally applied tension. It also reproduces the expected relations between internal and external stresses ^84^ (see Supplementary Fig. 17).

#### 2. Constant Frame Area Ensemble (*N, A*_*P*_, *T*)

In this ensemble the frame area (projected area), *A*_*P*_, is fixed. No additional frame tension term − *τA*_*P*_ is applied. This setup is useful for increased sampling at a constant frame area to isolate effects at fixed stages of membrane stretching on necks without externally applied tension.

### Generating configurations for high-genus vesicles and flat double membranes with PBC

Stomatocyte structures for this setup were obtained by relaxing and equilibrating a cuboid membrane configuration with *n* necks for genus *n* (for details see Supplementary Fig. 18). An expanding hard core potential was then placed at the centre of the shape, with an initial exclusion radius of 5*l*_*DTS*_. This radius was chosen to fit inside the innermost compartment of the stomatocyte. Expanding core simulations were performed over 5 × 10^6^ MC steps, with three replica each. The hard core potential was set to continually expand at 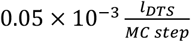. Since no surface vertex is allowed to lie inside the spherical core, the membrane must expand the centre compartment accordingly. When the bead radius increases, it is possible that some membrane vertices may initially lie inside the bead. However, after a few MC moves, they will all leave the volume of the bead, since only moves taking them outside the bead will be accepted. No additional volume control was applied. The surface area was effectively fixed with area compressibility *K*_*A*_ = 1000 *k*_*B*_*T* (see formula (5)). From these simulations, frames were taken at different sizes of core radius and equilibrated at that fixed radius over 3 − 5 × 10^6^ MC steps. Equilibrated morphologies were analysed for neck diameters and IM-OM distance. Details on their calculation can be found below.

Configurations for high-genus surfaces in PBC were obtained by relaxing and equilibrating a cuboid membrane configuration with open sides with one to six necks (genus 1-6) in periodic boundary conditions (for details see Supplementary Fig. 18). The simulations incorporate membrane fluctuations and allow for variable neck shapes dictated purely by free energy optimisation. To include the effect of mechanical stresses present during exponential cell growth or differentiation, which are assumed to act as tension on the NE membrane ^31,32,60^, we subjected our membrane to lateral tension with parameter τ (in the x-y-plane) and, taking into account Helfrich bending energy, determined its minimal free energy state.

For simulations over a range of frame areas, individual frames (states) were chosen from a simulation trajectory under tension. They were then equilibrated at constant frame area (*A*_*P*_)_*i*_ for the *i*-th frame, over 2 × 10^6^ MC steps. This was performed with a potential promoting constant surface area for area compressibilities of *K*_*A*_ = 0, 10, 100, 1000 *k*_*B*_*T* (see formula (5)). Equilibrated morphologies were analysed for bending energy, neck diameters, IM-OM distance and surface area at all frame areas (*A*_*P*_)_*i*._

### Analysis

For high-genus vesicles, neck diameter distributions and IM-OM distances were obtained from the last 50 frames of equilibrated simulations. At each frame, corresponding to one membrane mesh configuration, we first constructed the convex hull of this mesh. The mesh was then sectioned into IM and OM by distance of vertices to the convex hull, as the OM naturally lies very close to the hull, while the IM does not (see Fig. 6a, left). The threshold distance below (above) which a vertex was assigned to the OM (IM) was determined for each mesh individually, as it is influenced by the shape of each mesh and the IM-OM distance. The average IM-OM distance for each membrane configuration was subsequently determined by calculating the difference in the average distance of IM and OM to the vesicle centre (Fig. 6a, middle). Neck diameters *D*_*P*_ were calculated from the IM (or OM) through determining the circumference C of the necks, which were cut open due to separation of IM and OM mesh (see Fig. 6a, right).

**Fig. 6:**
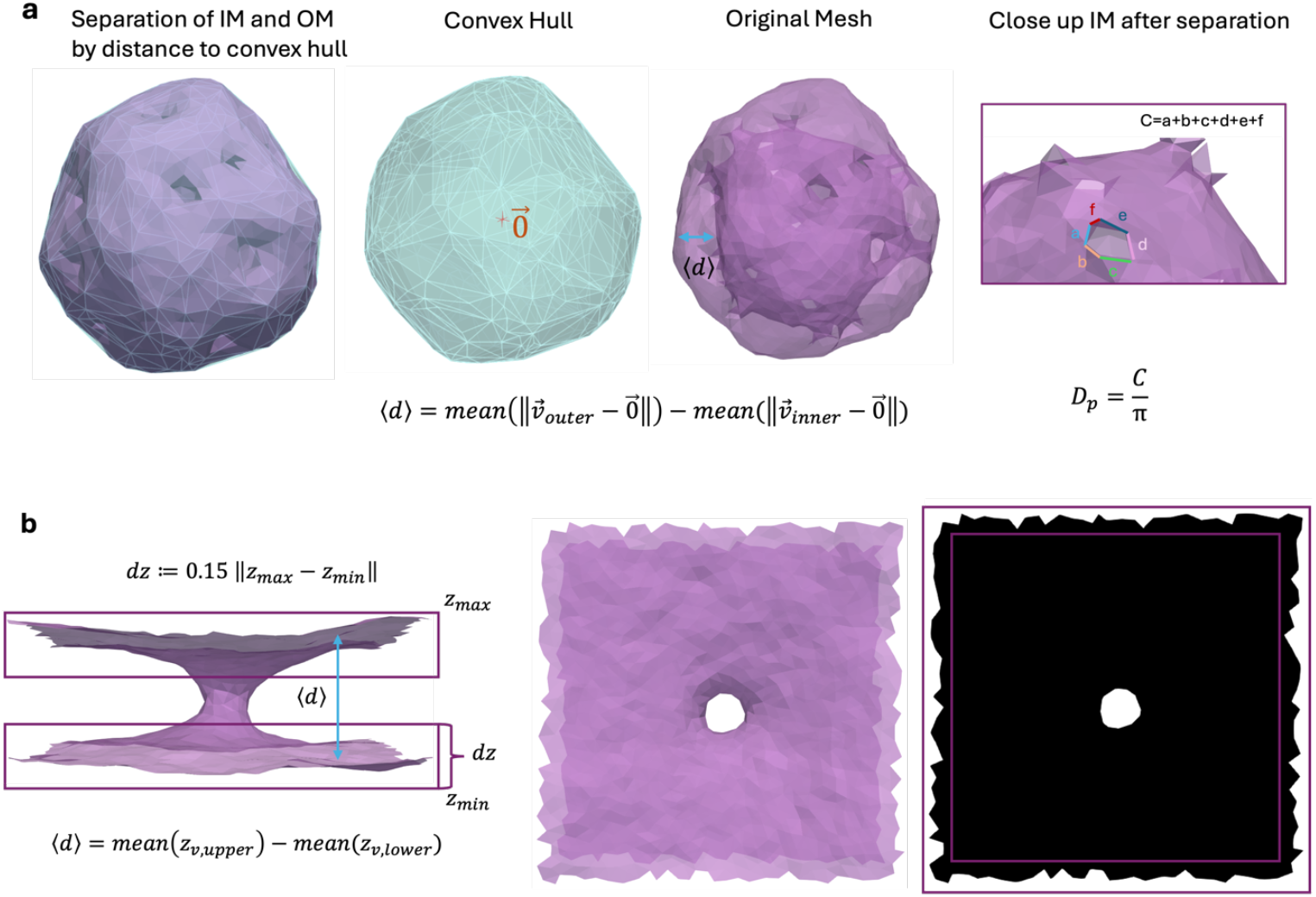
Process of determining IM-OM distance and membrane neck diameters. **(a)** For high-genus vesicle membranes, the convex hull was used as a criterion to divide the mesh into IM and OM. Vertices closer to the hull than a threshold value were assigned to the OM. Based on this separation into two meshes, membrane distance and neck diameters were determined. **(b)** For membranes in PBC, IM and OM were determined by the z-coordinate of vertices on the mesh as shown. Neck diameters were calculated based on a 2d-projection of the mesh.

For membranes in PBC, the average IM-OM distance was obtained by determining the average z-position of the two membrane sheets in the box and then calculating their distance (Fig. 6b, left). The neck diameter *D*_*P*_ was determined by projecting the mesh onto the x-y-plane and converting it to a binary image (Fig. 6b, middle & right). From the binary image, the area of the hole region *A*_*H*_ was determined by converting pixels to the simulation lengthscale using the boxsize known from the simulation. Then 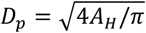. For extremely small neck sizes, this can underestimate neck size.

Every simulation in PBC was run with 10 replicas. Each datapoint for IM-OM distances, neck diameters and bending energies hence corresponds to the mean of the values recovered for these 10 replicas and errorbars indicate the standard error of the mean. Around branching points in frame area some replicas show high, others lower bending energy, indicating multiple local minima or degeneracy. When replica results are displayed individually instead of mean values, an error on the energy of each replica was obtained by block averaging over the last 10^6^ MC steps after equilibration of each simulation (20 blocks, length of 500 energy outputs; energy was saved every 100 MC steps). Only in case of genus-6, due to the larger system size, 8 × 10^5^ MC steps after equilibration were used for the analysis (16 blocks, length of 500 energy outputs; energy was saved every 100 MC steps).

### Proteins

Proteins in FreeDTS are modelled as inclusions with an in-plane orientation, so that there is at most one inclusion per membrane vertex ^7^. To emulate protein complexes that assemble in membrane necks, we chose proteins with asymmetric interactions with the membrane (inclusion type 2 in FreeDTS),

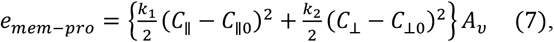

where *C*_∥_ (*C*_⊥_) denote membrane curvature parallel (perpendicular) to the inclusion orientation and *k*_1_, *k*_2_ are directional bending rigidities that the protein enforces on the membrane. Protein– protein interactions are modelled as a function of the angle between two inclusions *i, j* along the geodesic Θ = Θ_j_ − Θ_i_,

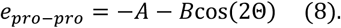

For more details see ^7^.

Stomatocytes were decorated by 10% of inclusions with 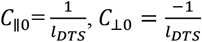, *k*_1_ = 10*k*_*B*_*T, k*_2_ = 5*k*_*B*_*T* and zero protein-protein interactions as this was enough for the proteins to cluster around the necks. Single membrane necks in PBC were decorated by 2.5-5% of inclusions with *C*_∥0_ in the range of 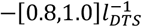, *k*_1_ = 10*k*_*B*_*T*. In these systems, attractive inclusion-inclusion interactions were used, *A* = 1*k*_*B*_*T* and *B* = (0,1,2,4)*k*_*B*_*T*, to explore larger parameter sets.

## Author Contributions

The manuscript was written through contributions of all authors. All authors have given approval to the final version of the manuscript. WP conceived the original idea. BG performed simulations, obtained the results and developed the theory. BG and WP wrote the manuscript. All authors discussed the results and commented on the manuscript at all stages.

## Notes

The authors declare no competing interests.

## ACKNOWLEDGMENT

We thank M Wood for comments and feedback on this manuscript.

## SUPPORTING INFORMATION AVAILABLE

Supplementary figures and notes with derivations and additional simulation details (Supplementary_Information.pdf).

Detailed tutorials, input files and code for data analysis publicly available on Github (https://github.com/weria-pezeshkian/FreeDTS/wiki/High-Genus--Membranes).

Supplementary movie of a membrane neck moving throughout a simulation in periodic boundary conditions (SmallNeck_Mobility_movie.avi).

## For Table of Contents Only

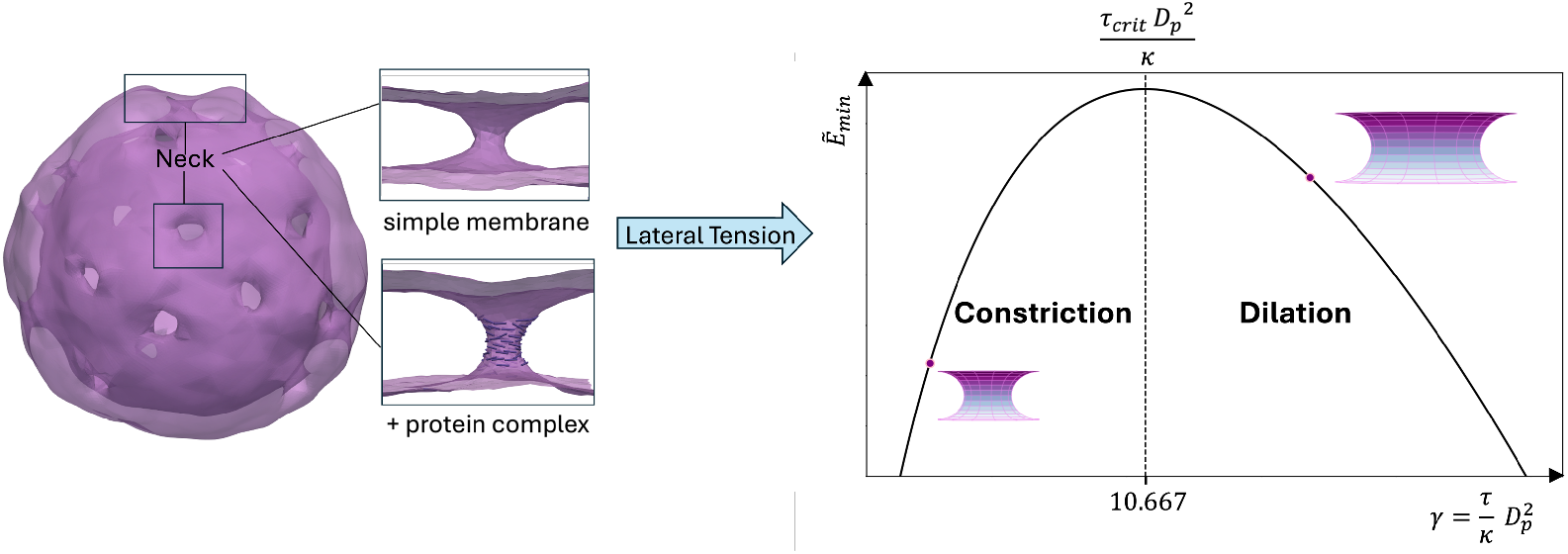

## REFERENCES

1. Gallagher, P. G. Chapter 45 Red Blood Cell Membrane Disorders. In Hematology (Seventh Edition); Hoffman, R., Benz, E. J., Silberstein, L. E., Heslop, H. E., Weitz, J. I., Anastasi, J., Salama, M. E., Abutalib, S. A., Eds.; Elsevier, 2018; pp 626–647. 10.1016/B978-0-323-35762-3.00045-7.

2. Lipowsky, R. The Conformation of Membranes. Nature 1991, 349 (6309), 475–481. 10.1038/349475a0.

3. Seifert, U.; Berndl, K.; Lipowsky, R. Shape Transformations of Vesicles: Phase Diagram for Spontaneous-Curvature and Bilayer-Coupling Models. Phys. Rev. A 1991, 44 (2), 1182– 1202. 10.1103/PhysRevA.44.1182.

4. De Franceschi, N.; Pezeshkian, W.; Fragasso, A.; Bruininks, B. M. H.; Tsai, S.; Marrink, S. J.; Dekker, C. Synthetic Membrane Shaper for Controlled Liposome Deformation. ACS Nano 2023, 17 (2), 966–978. 10.1021/acsnano.2c06125.

5. Lorent, J. H.; Levental, K. R.; Ganesan, L.; Rivera-Longsworth, G.; Sezgin, E.; Doktorova, M.; Lyman, E.; Levental, I. Plasma Membranes Are Asymmetric in Lipid Unsaturation, Packing and Protein Shape. Nat. Chem. Biol. 2020, 16 (6), 644–652. 10.1038/s41589-020-0529-6.

6. Pezeshkian, W.; König, M.; Marrink, S. J.; Ipsen, J. H. A Multi-Scale Approach to Membrane Remodeling Processes. Front. Mol. Biosci. 2019, 6. 10.3389/fmolb.2019.00059.

7. Pezeshkian, W.; Ipsen, J. H. Mesoscale Simulation of Biomembranes with FreeDTS. Nat. Commun. 2024, 15 (1), 548. 10.1038/s41467-024-44819-w.

8. Jülicher, F.; Seifert, U.; Lipowsky, R. Conformal Degeneracy and Conformal Diffusion of Vesicles. Phys. Rev. Lett. 1993, 71 (3), 452–455. 10.1103/PhysRevLett.71.452.

9. Noguchi, H. Construction of Nuclear Envelope Shape by a High-Genus Vesicle with Pore-Size Constraint. Biophys. J. 2016, 111 (4), 824–831. 10.1016/j.bpj.2016.07.010.

10. Noguchi, H. Shape Transitions of High-Genus Fluid Vesicles. Europhys. Lett. 2015, 112 (5), 58004. 10.1209/0295-5075/112/58004.

11. Otsuka, S.; Steyer, A. M.; Schorb, M.; Hériché, J.-K.; Hossain, M. J.; Sethi, S.; Kueblbeck, M.; Schwab, Y.; Beck, M.; Ellenberg, J. Postmitotic Nuclear Pore Assembly Proceeds by Radial Dilation of Small Membrane Openings. Nat. Struct. Mol. Biol. 2018, 25 (1), 21–28. 10.1038/s41594-017-0001-9.

12. Parzych, K. R.; Klionsky, D. J. An Overview of Autophagy: Morphology, Mechanism, and Regulation. Antioxid. Redox Signal. 2014, 20 (3), 460–473. 10.1089/ars.2013.5371.

13. De Magistris, P.; Antonin, W. The Dynamic Nature of the Nuclear Envelope. Curr. Biol. 2018, 28 (8), R487–R497. 10.1016/j.cub.2018.01.073.

14. Bieber, A.; Capitanio, C.; Erdmann, P. S.; Fiedler, F.; Beck, F.; Lee, C.-W.; Li, D.; Hummer, G.; Schulman, B. A.; Baumeister, W.; Wilfling, F. In Situ Structural Analysis Reveals Membrane Shape Transitions during Autophagosome Formation. Proc. Natl. Acad. Sci. U. S. A. 2022, 119 (39), e2209823119. 10.1073/pnas.2209823119.

15. Sakai, Y.; Takahashi, S.; Koyama-Honda, I.; Saito, C.; Mizushima, N. Experimental Determination and Mathematical Modeling of Standard Shapes of Forming Autophagosomes. Nat. Commun. 2024, 15 (1), 91. 10.1038/s41467-023-44442-1.

16. Sakai, Y.; Koyama-Honda, I.; Tachikawa, M.; Knorr, R. L.; Mizushima, N. Modeling Membrane Morphological Change during Autophagosome Formation. iScience 2020, 23 (9), 101466. 10.1016/j.isci.2020.101466.

17. Hetzer, M. W. The Nuclear Envelope. Cold Spring Harb. Perspect. Biol. 2010, 2 (3), a000539. 10.1101/cshperspect.a000539.

18. Webster, M.; Witkin, K. L.; Cohen-Fix, O. Sizing up the Nucleus: Nuclear Shape, Size and Nuclear-Envelope Assembly. J. Cell Sci. 2009, 122 (10), 1477–1486. 10.1242/jcs.037333.

19. Strambio-De-Castillia, C.; Niepel, M.; Rout, M. P. The Nuclear Pore Complex: Bridging Nuclear Transport and Gene Regulation. Nat. Rev. Mol. Cell Biol. 2010, 11 (7), 490–501. 10.1038/nrm2928.

20. Fabre, E.; Hurt, E. Yeast Genetics to Dissect the Nuclear Pore Complex and Nucleocytoplasmic Trafficking. Annu. Rev. Genet. 1997, 31 (Volume 31, 1997), 277–313. 10.1146/annurev.genet.31.1.277.

21. Grossman, E.; Medalia, O.; Zwerger, M. Functional Architecture of the Nuclear Pore Complex. Annu. Rev. Biophys. 2012, 41 (Volume 41, 2012), 557–584. 10.1146/annurev-biophys-050511-102328.

22. Hoelz, A.; Debler, E. W.; Blobel, G. The Structure of the Nuclear Pore Complex. Annu. Rev. Biochem. 2011, 80 (Volume 80, 2011), 613–643. 10.1146/annurev-biochem-060109-151030.

23. Patel, M. K.; Chakrabarti, B.; Panwar, A. S. Emergence of Selectivity and Specificity in a Coarse-Grained Model of the Nuclear Pore Complex with Sequence-Agnostic FG-Nups. Phys. Chem. Chem. Phys. 2023, 25 (48), 32824–32836. 10.1039/D3CP03746K.

24. Efremov, A. K.; Hovan, L.; Yan, J. Nucleus Size and Its Effect on Nucleosome Stability in Living Cells. Biophys. J. 2022, 121 (21), 4189–4204. 10.1016/j.bpj.2022.09.019.

25. Stephens, A. D.; Banigan, E. J.; Adam, S. A.; Goldman, R. D.; Marko, J. F. Chromatin and Lamin A Determine Two Different Mechanical Response Regimes of the Cell Nucleus. Mol. Biol. Cell 2017, 28 (14), 1984–1996. 10.1091/mbc.e16-09-0653.

26. Mazumder, A.; Roopa, T.; Basu, A.; Mahadevan, L.; Shivashankar, G. V. Dynamics of Chromatin Decondensation Reveals the Structural Integrity of a Mechanically Prestressed Nucleus. Biophys. J. 2008, 95 (6), 3028–3035. 10.1529/biophysj.108.132274.

27. Finan, J. D.; Chalut, K. J.; Wax, A.; Guilak, F. Nonlinear Osmotic Properties of the Cell Nucleus. Ann. Biomed. Eng. 2009, 37 (3), 477–491. 10.1007/s10439-008-9618-5.

28. Dahl, K. N.; Kahn, S. M.; Wilson, K. L.; Discher, D. E. The Nuclear Envelope Lamina Network Has Elasticity and a Compressibility Limit Suggestive of a Molecular Shock Absorber. J. Cell Sci. 2004, 117 (20), 4779–4786. 10.1242/jcs.01357.

29. Deviri, D.; Safran, S. A. Balance of Osmotic Pressures Determines the Volume of the Cell Nucleus. bioRxiv October 1, 2021, p 2021.10.01.462771. 10.1101/2021.10.01.462771.

30. Lemière, J.; Real-Calderon, P.; Holt, L. J.; Fai, T. G.; Chang, F. Control of Nuclear Size by Osmotic Forces in Schizosaccharomyces Pombe. eLife 2022, 11, e76075. 10.7554/eLife.76075.

31. Zimmerli, C. E.; Allegretti, M.; Rantos, V.; Goetz, S. K.; Obarska-Kosinska, A.; Zagoriy, I.; Halavatyi, A.; Hummer, G.; Mahamid, J.; Kosinski, J.; Beck, M. Nuclear Pores Dilate and Constrict in Cellulo. Science 2021, 374 (6573), eabd9776. 10.1126/science.abd9776.

32. Taniguchi, R.; Orniacki, C.; Kreysing, J. P.; Zila, V.; Zimmerli, C. E.; Böhm, S.; Turoňová, B.; Kräusslich, H.-G.; Doye, V.; Beck, M. Nuclear Pores Safeguard the Integrity of the Nuclear Envelope. Nat. Cell Biol. 2025, 1–14. 10.1038/s41556-025-01648-3.

33. Schuller, A. P.; Wojtynek, M.; Mankus, D.; Tatli, M.; Kronenberg-Tenga, R.; Regmi, S. G.; Dip, P. V.; Lytton-Jean, A. K. R.; Brignole, E. J.; Dasso, M.; Weis, K.; Medalia, O.; Schwartz, T. U. The Cellular Environment Shapes the Nuclear Pore Complex Architecture. Nature 2021, 598 (7882), 667–671. 10.1038/s41586-021-03985-3.

34. Mosalaganti, S.; Obarska-Kosinska, A.; Siggel, M.; Taniguchi, R.; Turoňová, B.; Zimmerli, C. E.; Buczak, K.; Schmidt, F. H.; Margiotta, E.; Mackmull, M.-T.; Hagen, W. J. H.; Hummer, G.; Kosinski, J.; Beck, M. AI-Based Structure Prediction Empowers Integrative Structural Analysis of Human Nuclear Pores. Science 2022, 376 (6598), eabm9506. 10.1126/science.abm9506.

35. Hoffmann, P. C.; Kim, H.; Obarska-Kosinska, A.; Kreysing, J. P.; Andino-Frydman, E.; Cruz-León, S.; Margiotta, E.; Cernikova, L.; Kosinski, J.; Turoňová, B.; Hummer, G.; Beck, M. Nuclear Pore Permeability and Fluid Flow Are Modulated by Its Dilation State. Mol. Cell 2025, 85 (3), 537-554.e11. 10.1016/j.molcel.2024.11.038.

36. Bui, K. H.; von Appen, A.; DiGuilio, A. L.; Ori, A.; Sparks, L.; Mackmull, M.-T.; Bock, T.; Hagen, W.; Andrés-Pons, A.; Glavy, J. S.; Beck, M. Integrated Structural Analysis of the Human Nuclear Pore Complex Scaffold. Cell 2013, 155 (6), 1233–1243. 10.1016/j.cell.2013.10.055.

37. Yu, M.; Heidari, M.; Mikhaleva, S.; Tan, P. S.; Mingu, S.; Ruan, H.; Reinkemeier, C. D.; Obarska-Kosinska, A.; Siggel, M.; Beck, M.; Hummer, G.; Lemke, E. A. Visualizing the Disordered Nuclear Transport Machinery in Situ. Nature 2023, 617 (7959), 162–169. 10.1038/s41586-023-05990-0.

38. Bottacchiari, M.; Gallo, M.; Bussoletti, M.; Casciola, C. M. Activation Energy and Force Fields during Topological Transitions of Fluid Lipid Vesicles. Commun. Phys. 2022, 5 (1), 283. 10.1038/s42005-022-01055-2.

39. Knecht, V.; Marrink, S.-J. Molecular Dynamics Simulations of Lipid Vesicle Fusion in Atomic Detail. Biophys. J. 2007, 92 (12), 4254–4261. 10.1529/biophysj.106.103572.

40. Kawamoto, S.; Shinoda, W. Free Energy Analysis along the Stalk Mechanism of Membrane Fusion. Soft Matter 2014, 10 (17), 3048–3054. 10.1039/C3SM52344F.

41. Kawamoto, S.; Klein, M. L.; Shinoda, W. Coarse-Grained Molecular Dynamics Study of Membrane Fusion: Curvature Effects on Free Energy Barriers along the Stalk Mechanism. J. Chem. Phys. 2015, 143 (24), 243112. 10.1063/1.4933087.

42. Roffay, C.; Molinard, G.; Kim, K.; Urbanska, M.; Andrade, V.; Barbarasa, V.; Nowak, P.; Mercier, V.; García-Calvo, J.; Matile, S.; Loewith, R.; Echard, A.; Guck, J.; Lenz, M.; Roux, Passive Coupling of Membrane Tension and Cell Volume during Active Response of Cells to Osmosis. Proc. Natl. Acad. Sci. 2021, 118 (47), e2103228118. 10.1073/pnas.2103228118.

43. Alam Shibly, S. U.; Ghatak, C.; Sayem Karal, M. A.; Moniruzzaman, Md.; Yamazaki, M. Experimental Estimation of Membrane Tension Induced by Osmotic Pressure. Biophys. J. 2016, 111 (10), 2190–2201. 10.1016/j.bpj.2016.09.043.

44. Lammerding, J. Mechanics of the Nucleus. Compr. Physiol. 2011, 1 (2), 783–807. 10.1002/cphy.c100038.

45. Pezeshkian, W.; Ipsen, J. H. Fluctuations and Conformational Stability of a Membrane Patch with Curvature Inducing Inclusions. Soft Matter 2019, 15 (48), 9974–9981. 10.1039/C9SM01762C.

46. Neder, J.; West, B.; Nielaba, P.; Schmid, F. Coarse-Grained Simulations of Membranes under Tension. J. Chem. Phys. 2010, 132 (11), 115101. 10.1063/1.3352583.

47. Shiba, H.; Noguchi, H.; Fournier, J.-B. Monte Carlo Study of the Frame, Fluctuation and Internal Tensions of Fluctuating Membranes with Fixed Area. Soft Matter 2016, 12 (8), 2373–2380. 10.1039/C5SM01900A.

48. Durand, M. Frame Tension Governs the Thermal Fluctuations of a Fluid Membrane: New Evidence. Soft Matter 2022, 18 (20), 3891–3901. 10.1039/D1SM01765A.

49. Maul, G.; Deaven, L. Quantitative Determination of Nuclear Pore Complexes in Cycling Cells with Differing DNA Content. J. Cell Biol. 1977, 73 (3), 748–760.

50. Chizmadzhev, Y. A.; Cohen, F. S.; Shcherbakov, A.; Zimmerberg, J. Membrane Mechanics Can Account for Fusion Pore Dilation in Stages. Biophys. J. 1995, 69 (6), 2489–2500. 10.1016/S0006-3495(95)80119-0.

51. Jackson, M. B. Minimum Membrane Bending Energies of Fusion Pores. J. Membr. Biol. 2009, 231 (2–3), 101–115. 10.1007/s00232-009-9209-x.

52. Lin, D. H.; Hoelz, A. The Structure of the Nuclear Pore Complex (An Update). Annu. Rev. Biochem. 2019, 88, 725–783. 10.1146/annurev-biochem-062917-011901.

53. Bochicchio, D.; Monticelli, L. Chapter Five - The Membrane Bending Modulus in Experiments and Simulations: A Puzzling Picture. In Advances in Biomembranes and Lipid Self-Assembly; Iglič, A., Kulkarni, C. V., Rappolt, M., Eds.; Academic Press, 2016; Vol. 23, pp 117–143. 10.1016/bs.abl.2016.01.003.

54. Torbati, M.; Lele, T. P.; Agrawal, A. Ultradonut Topology of the Nuclear Envelope. Proc. Natl. Acad. Sci. U. S. A. 2016, 113 (40), 11094–11099. 10.1073/pnas.1604777113.

55. Gauthier, N. C.; Masters, T. A.; Sheetz, M. P. Mechanical Feedback between Membrane Tension and Dynamics. Trends Cell Biol. 2012, 22 (10), 527–535. 10.1016/j.tcb.2012.07.005.

56. Agrawal, A.; Lele, T. P. Mechanics of Nuclear Membranes. J. Cell Sci. 2019, 132 (14), jcs229245. 10.1242/jcs.229245.

57. Michalet, X.; Bensimon, D.; Fourcade, B. Fluctuating Vesicles of Nonspherical Topology. Phys. Rev. Lett. 1994, 72 (1), 168–171. 10.1103/PhysRevLett.72.168.

58. Jülicher, F. The Morphology of Vesicles of Higher Topological Genus: Conformal Degeneracy and Conformal Modes. J. Phys. II 1996, 6 (12), 1797–1824. 10.1051/jp2:1996161.

59. Frolov, V. A.; Lizunov, V. A.; Dunina-Barkovskaya, A. Ya.; Samsonov, A. V.; Zimmerberg, J. Shape Bistability of a Membrane Neck: A Toggle Switch to Control Vesicle Content Release. Proc. Natl. Acad. Sci. 2003, 100 (15), 8698–8703. 10.1073/pnas.1432962100.

60. Matsuda, A.; Mofrad, M. R. K. On the Nuclear Pore Complex and Its Emerging Role in Cellular Mechanotransduction. APL Bioeng. 2022, 6 (1), 011504. 10.1063/5.0080480.

61. Schirmer, E. C.; Florens, L.; Guan, T.; Yates, J. R.; Gerace, L. Nuclear Membrane Proteins with Potential Disease Links Found by Subtractive Proteomics. Science 2003, 301 (5638), 1380–1382. 10.1126/science.1088176.

62. Akhtar, A.; Gasser, S. M. The Nuclear Envelope and Transcriptional Control. Nat. Rev. Genet. 2007, 8 (7), 507–517. 10.1038/nrg2122.

63. Dorner, D.; Gotzmann, J.; Foisner, R. Nucleoplasmic Lamins and Their Interaction Partners, LAP2α, Rb, and BAF, in Transcriptional Regulation. FEBS J. 2007, 274 (6), 1362–1373. 10.1111/j.1742-4658.2007.05695.x.

64. Beiter, J.; Voth, G. A. Making the Cut: Multiscale Simulation of Membrane Remodeling. Curr. Opin. Struct. Biol. 2024, 87, 102831. 10.1016/j.sbi.2024.102831.

65. Pezeshkian, W.; Marrink, S. J. Simulating Realistic Membrane Shapes. Curr. Opin. Cell Biol. 2021, 71, 103–111. 10.1016/j.ceb.2021.02.009.

66. Evans, E.; Heinrich, V.; Ludwig, F.; Rawicz, W. Dynamic Tension Spectroscopy and Strength of Biomembranes. Biophys. J. 2003, 85 (4), 2342–2350. 10.1016/S0006-3495(03)74658-X.

67. Evans, E.; Smith, B. A. Kinetics of Hole Nucleation in Biomembrane Rupture. New J. Phys. 2011, 13 (9), 095010. 10.1088/1367-2630/13/9/095010.

68. Lipowsky, R.; Pramanik, S.; Benk, A. S.; Tarnawski, M.; Spatz, J. P.; Dimova, R. Elucidating the Morphology of the Endoplasmic Reticulum: Puzzles and Perspectives. ACS Nano 2023, 17 (13), 11957–11968. 10.1021/acsnano.3c01338.

69. Upadhyaya, A.; Sheetz, M. P. Tension in Tubulovesicular Networks of Golgi and Endoplasmic Reticulum Membranes. Biophys. J. 2004, 86 (5), 2923–2928. 10.1016/S0006-3495(04)74343-X.

70. Zucker, B.; Golani, G.; Kozlov, M. M. Model for Ring Closure in ER Tubular Network Dynamics. Biophys. J. 2023, 122 (11), 1974–1984. 10.1016/j.bpj.2022.10.005.

71. Otsuka, S.; Steyer, A. M.; Schorb, M.; Hériché, J.-K.; Hossain, M. J.; Sethi, S.; Kueblbeck, M.; Schwab, Y.; Beck, M.; Ellenberg, J. Postmitotic Nuclear Pore Assembly Proceeds by Radial Dilation of Small Membrane Openings. Nat. Struct. Mol. Biol. 2018, 25 (1), 21–28. 10.1038/s41594-017-0001-9.

72. Kroll, D. M.; Gompper, G. The Conformation of Fluid Membranes: Monte Carlo Simulations. Science 1992, 255 (5047), 968–971. 10.1126/science.1546294.

73. Dasanna, A. K.; Fedosov, D. A. Mesoscopic Modeling of Membranes at Cellular Scale. Eur. Phys. J. Spec. Top. 2024, 233 (21), 3053–3071. 10.1140/epjs/s11734-024-01177-4.

74. Duncan, A. L.; Pezeshkian, W. Mesoscale Simulations: An Indispensable Approach to Understand Biomembranes. Biophys. J. 2023, 122 (11), 1883–1889. 10.1016/j.bpj.2023.02.017.

75. Siggel, M.; Kehl, S.; Reuter, K.; Köfinger, J.; Hummer, G. TriMem: A Parallelized Hybrid Monte Carlo Software for Efficient Simulations of Lipid Membranes. J. Chem. Phys. 2022, 157 (17), 174801. 10.1063/5.0101118.

76. Harker-Kirschneck, L.; Hafner, A. E.; Yao, T.; Vanhille-Campos, C.; Jiang, X.; Pulschen, A.; Hurtig, F.; Hryniuk, D.; Culley, S.; Henriques, R.; Baum, B.; Šarić, A. Physical Mechanisms of ESCRT-III–Driven Cell Division. Proc. Natl. Acad. Sci. 2022, 119 (1), e2107763119. 10.1073/pnas.2107763119.

77. Sadeghi, M.; Noé, F. Large-Scale Simulation of Biomembranes Incorporating Realistic Kinetics into Coarse-Grained Models. Nat. Commun. 2020, 11 (1), 2951. 10.1038/s41467-020-16424-0.

78. Davtyan, A.; Simunovic, M.; Voth, G. A. The Mesoscopic Membrane with Proteins (MesM-P) Model. J. Chem. Phys. 2017, 147 (4), 044101. 10.1063/1.4993514.

79. Ayton, G. S.; Blood, P. D.; Voth, G. A. Membrane Remodeling from N-BAR Domain Interactions: Insights from Multi-Scale Simulation. Biophys. J. 2007, 92 (10), 3595–3602. 10.1529/biophysj.106.101709.

80. Pezeshkian, W.; Ipsen, J. H. Creasing of Flexible Membranes at Vanishing Tension. Phys. Rev. E 2021, 103 (4), L041001. 10.1103/PhysRevE.103.L041001.

81. Helfrich, W.; Servuss, R.-M. Undulations, Steric Interaction and Cohesion of Fluid Membranes. Il Nuovo Cimento D 1984, 3 (1), 137–151. 10.1007/BF02452208.

82. Goetz, R.; Gompper, G.; Lipowsky, R. Mobility and Elasticity of Self-Assembled Membranes. Phys. Rev. Lett. 1999, 82 (1), 221–224. 10.1103/PhysRevLett.82.221.

83. Venturoli, M.; Sperotto, M. M.; Kranenburg, M.; Smit, B. xMesoscopic Models of Biological Membranes. Phys. Rep. 2006, 437. 10.1016/j.physrep.2006.07.006.

84. Lipowsky, R. The Many Faces of Membrane Tension for Biomembranes and Vesicles. Faraday Discuss. 2025, 259 (0), 234–263. 10.1039/D4FD00184B.

