## Supplementary Notes and Figures for "Bimodal Mechanical Response of Membrane Necks: Implications for the Nuclear Envelope"

#### Supplementary Note 1

##### 1.1 Osmotic Pressure in the IM-OM lumen volume

In the following we show that the effect of an osmotic pressure on the lumen volume between inner and outer membrane (Compt. II) of a high-genus stomatocyte, like the nuclear envelope (NE), should be significantly smaller than the effect on the internal volume (Compt. I). The main reason is that the lumen volume is significantly smaller than the internal volume. Therefore, in modelling the effect of osmotic pressure here, it is reasonable to assume that the internal volume will be mostly controlled, while a lumen volume constraint plays a subordinate role or is even negligible.

To show this, we consider the Jacobus van 't Hoff equation for osmotic pressure, $\Pi = icRT$, where $i$ is the van 't Hoff index, $c$ is the molar concentration of solute, $R$ is the ideal gas constant, and $T$ is the temperature. For a vesicle system, let $V_{ini}$ be the initial vesicle volume and $\bar{c}_j = \sum_n i_n c_n$ the effective compartment concentration of solute. Here, $j$ is either inside the compartment (as $V_{ini}$), or outside. The summation includes all solute types $n$ in the compartment $j$. Then $\Delta\Pi = RT\left(\frac{V_{ini}\bar{c}_{in}}{V} - \bar{c}_{out}\right)$.

Therefore, the energy associated with changes of the vesicle volume from $V_{ini}$ to $V$ will be:

$$\Delta E_{osmos}(V) = -RT\int_{V_{ini}}^{V} \Delta\Pi dV = -RT\left[\bar{c}_{in}V_{ini}ln\left(\frac{V}{V_{ini}}\right) - \bar{c}_{out}(V - V_{ini})\right] \quad (2)$$

$$= RT\bar{c}_{out}V_{eq}\left(\frac{1}{2}\left(\frac{V-V_{eq}}{V_{eq}}\right)^2 + \left[\sum_{i=3}^{\infty}\left(\frac{V_{eq}-V}{iV_{eq}}\right)^i\right]\right) \quad (3).$$

where $\bar{c}_{out}V_{eq} = \bar{c}_{in}V_{ini}$ and, importantly $K \coloneqq RT\bar{c}_{out}V_{eq}$, so the compartments adapt their volume to $V = V_{eq} = \frac{\bar{c}_{in}V_{ini}}{\bar{c}_{out}}$. The coupling constant $K$ for the lumen volume will be significantly smaller than the one for the internal volume, as it scales with the respective compartment volume (as the ion concentration are in the same order of magnitude, order of $100\,mM$). Hence, an ion concentration difference will only weakly influence the lumen volume compared to the internal volume. This can be seen as follows:
Assuming a spherical NE, let $R$ be the radius of the internal compartment; then the internal volume is $V_i = \frac{4}{3}\pi R^3$, while the lumen volume is $V_l = \frac{4}{3}\pi(3hR^2 + 3h^2R + h^3)$, $h$ being the distance between the two membranes.

The coupling constants for the volume of internal part ($K_i$) vs. lumen ($K_l$) then scale as:

$$\frac{K_l}{K_i} \approx \frac{3h}{R} \approx 10^{-2} - 10^{-3},$$

since for the NE one has $h \approx 20nm$ [1], while $R \approx 2.5 - 10\mu m$ [2].

##### 1.2 Estimating nuclear pore distances

In human cells the nuclear envelope has an outer surface of $\approx 300\ \mu m^2$ with a nuclear pore density of $8 - 12\ /\mu m^2$ [3]. To roughly estimate a minimum average distance between nuclear pores, we assume each NPC has an area of $\frac{1}{12}\mu m^2 \approx 0.083\ \mu m^2$ available within which no other nuclear pores reside. Assuming a square (or circular) shape for this area leads to an estimated distance of $\frac{\sqrt{0.083\ \mu m^2}}{2} \approx 150\ nm$ ($\sqrt{0.083\ \mu m^2/\pi} \approx 160\ nm$) between each two nuclear pores.

#### Supplementary Note 2

##### Ensemble of constant frame tension

Such a system can be described by a Helmholtz free energy, $F(A, A_p, A_0, \dots)$; where $A$ is surface area, $A_p$ is the projected area (box dimension in the XY plane), $A_0$ is the number of the molecules multiplied by the area per molecule (different representation of particle number). Please note that the three dots stand for other macroscopic variables such as temperature and bending rigidity and not membrane tensions. From the Helmholtz free energy, the tensions can in principle be determined as

$$\tau = \left(\frac{\partial F}{\partial A_p}\right)_{A_0, A, \dots} \text{ and } \sigma = \left(\frac{\partial F}{\partial A}\right)_{A_0, A_p, \dots} \quad (13).$$

Here, both tensions are dependent on model parameters and the constraints. For an ensemble in which $\tau$ is fixed (with variable $A_p$), the relevant Legendre transformed free energy (here denoted as $R$) will be $R(A, \tau, A_0, \dots) = F\big(A, A_p, A_0, \dots\big) - \tau A_p$ and $\sigma$ can be obtained as

$$\sigma = \left(\frac{\partial R}{\partial A}\right)_{A_0, \tau, \dots} \quad (14).$$

In particular, if the density of the necks is smaller than the surface area one can consider the rest of the surface to be parallel.

#### Supplementary Note 3

##### Theory: Toroidal membrane neck under lateral tension; constant surface area

To describe the toroidal membrane neck under tension, we consider the neck to be embedded in two square pieces of membrane with side length $L$, corresponding to inner and outer membrane (IM & OM). We assume that the neck is toroidal with diameters $D_p$ and $d$, defining the smallest part of the channel and half of the INM-ONM distance respectively, and that the other parts of the membrane are flat. We set up a total energy to minimise, to reveal the optimal dimension of the neck in terms of the two torus diameters, chosen as shown in the main text (Figure 1A), where $d$ defines the distance between the two membrane sheets, while $D_p$ spans the tightest part of the neck.

Our energy to minimise will be $E_{tot} = E_B + E_\tau$, considering the bending energy of the membrane, as well as the contribution from lateral tension. We furthermore assume a constant total surface area, $A_0$. The bending energy of the system can be described by the Helfrich Hamiltonian,

$$E_B = \oint \left[\frac{\kappa}{2}(2H)^2 - \kappa_G K\right] dA \quad (1),$$

with bending modulus $\kappa$, and gaussian modulus $\kappa_G$. For a toroidal neck, the principal curvatures in spherical coordinates are

$$c_1 = \frac{2}{d}, c_2 = \frac{2cos(\phi)}{D_p + d + dcos(\phi)} \quad (2).$$

By integrating over the surface of the toroidal neck, one finds a constant contribution of Gaussian curvature, which will hence be neglected going forward. The mean curvature contribution amounts to a bending energy of

$$E_B = f\left(\frac{D_p}{d}\right) = 4\pi\kappa \frac{D_p}{d} \frac{\left(1+\frac{d}{D_p}\right)^2}{\sqrt{1+\frac{2d}{D_p}}} tan^{-1}\left(\sqrt{\frac{2d}{D_p}+1}\right) \quad (3),$$

for the toroidal neck. Lateral tension is represented by an energetic contribution $E_\tau = -\tau A_p$ with tension parameter $\tau$ and projected area $A_p$. The constraint of constant surface area yields

$$A_p = \frac{A_0}{2} + \pi\frac{{D_p}^2}{4} + \frac{D_p d}{4}(2\pi - \pi^2) + \frac{d^2}{4}(3\pi - \pi^2) \quad (4).$$

Considering the bending and tension term separately, grants additional intuition about the system. The bending energy depends solely on the ratio of diameters,

$$E_B = f\left(\frac{D_p}{d}\right) = \tilde{f}\left(\frac{d}{D_p}\right) \quad (5),$$

and minimises at $\frac{D_p}{d} \approx 0.6$ (see Suppl. Fig. 8). In contrast, tension,

$$E_\tau = -\tau\pi\left[\frac{A_0}{2\pi} + \frac{{D_p}^2}{4} + \frac{D_p d}{4}(2-\pi) + \frac{d^2}{4}(3-\pi)\right] \quad (6)$$

decreases with any increase in projected area, which favors increasing $D_p$, but not $d$. Hence, the optimal diameters are determined by the interplay of bending and tension energy.

We then rescaled the energy to minimise it with respect to $s = \frac{d}{D_p}$ at constant values of the dimensionless variable, $\gamma = \frac{\tau}{\kappa}{D_p}^2$, that combines the effect of tension parameter, bending rigidity and neck diameter:

$$\tilde{E} = g(\gamma, s) = \frac{E_B + E_\tau}{4\pi\kappa} + \frac{\tau}{\kappa}\frac{A_0}{2\pi} = \frac{E_B(s)}{4\pi\kappa} + \frac{E_\tau(\gamma, s)}{4\pi\kappa} + \frac{\tau}{\kappa}\frac{A_0}{2\pi} \quad (7),$$

One can exploit the fact that for each fixed value of $\gamma$, the total energy has one unique minimum in $s$. Determining this minimum numerically over a fine grid of values for fixed $\gamma$, we recover the minimal energy as a function of $\gamma$-values, $\tilde{E}_{min}(\gamma)$, which reveals the interplay of the two energetic contributions of bending and tension and how they determine the neck size, together with the starting diameter of the neck (see main text Figure 3A).

We now incorporate the effect of a small spontaneous curvature $C_0$. Let $E_B$ denote the bending energy in the toroidal neck as before at $C_0 = 0$. Then:

$$E_{B,C_0} = \oint \frac{\kappa}{2}(2H - C_0)^2 dA = E_B - \oint 2\kappa H C_0 dA + \oint \frac{\kappa}{2} C_0^2 dA \ (8)$$

Using the principal curvatures of the toroidal neck (eq. (2)), this yields:

$$E_{B1,C_0} = E_B + 4\,\pi\kappa\left\{-\frac{C_0\pi}{4}\left[D_p + d\left(1 - \frac{4}{\pi}\right)\right] + \frac{C_0^2\pi}{16}\left[D_p d + d^2\left(1 - \frac{2}{\pi}\right)\right]\right\} (9).$$

To account for effect of $C_0$ on the flat membrane patches, another term must be taken into account. The area of the flat parts alone will be $A_f = A_p - \frac{(D_p + d)^2}{2}$. Using eq. (4) this will be:

$$E_{B2,C_0} = \frac{\kappa}{2} C_0^2 \pi\left[\frac{A_0}{2\pi} - \frac{D_p^2}{4} - \frac{D_p d}{4}(2+\pi) + \frac{d^2}{4}(1-\pi)\right] (10).$$

Before minimizing the total energy, $E_{B1,C_0} + E_{B2,C_0} + E_\tau$, we want to rescale it again as previously, using $s = \frac{d}{D_p}$ and $\gamma = \frac{\tau}{\kappa}{D_p}^2$. Neglecting constant terms and using eq. (5):

$$\begin{aligned} E_{B,C_0} &= E_{B1,C_0} + E_{B2,C_0} \\ &= \tilde{f}(s) + 4\,\pi\kappa\left\{\frac{C_0^2\pi}{16} D_p^2\left[s + s^2\left(1 - \frac{2}{\pi}\right)\right] - \frac{C_0\pi}{4} D_p\left[1 + s\left(1 - \frac{4}{\pi}\right)\right]\right\} \end{aligned}$$

$$-\frac{\pi\kappa C_0^2 D_p^2}{2}\left[\frac{1}{4}+\frac{s}{4}(2+\pi)-\frac{s^2}{4}(1-\pi)\right] \text{ (11).}$$

Then we can rescale as previously and arrive at:

$$\widetilde{E_{B,C_0}} = \frac{\tilde{f}(s)}{4\,\pi\kappa} + \left\{\gamma\frac{\kappa}{\tau}C_0^2\left[\frac{s}{32}(\pi-2)+\frac{s^2}{32}(\pi-3)-\frac{1}{32}\right]-\sqrt{\gamma}\sqrt{\frac{\kappa}{\tau}}C_0\left[\frac{\pi}{4}+\frac{s}{4}(\pi-4)\right]\right\} \text{ (12).}$$

Defining $\tilde{C} = \sqrt{\frac{\kappa}{\tau}}C_0$ and the previous expression for the total rescaled energy $\tilde{E}$ (also containing tension) becomes:

$$\widetilde{E_{C_0}} = \tilde{E} + \gamma\tilde{C}^2\left[\frac{s}{32}(\pi-2)+\frac{s^2}{32}(\pi-3)-\frac{1}{32}\right] - \sqrt{\gamma}\,\tilde{C}\left[\frac{\pi}{4}+\frac{s(\pi-4)}{4}\right] \text{ (13).}$$

Again, one can numerically minimize the total energy with respect to $s$ over a fine grid of $\gamma$-values and we recover the minimal energy as $\widetilde{E_{C_0}}_{min}(\gamma)$, but this time for $C_0 \neq 0$. Increasing spontaneous curvature shifts the maximum in $\widetilde{E_{C_0}}_{min}(\gamma)$ to smaller $\gamma$, thereby shortening the region of positive slope until it disappears for $\tilde{C} > 0.19$ (Suppl. Fig. 9A). Spontaneous curvature can furthermore cause emergence of a new local minimum at small $\gamma$ values. We find this minimum to be present for $0.0 < \tilde{C} \leq 0.2$. Suppl. Fig. 9B shows how the (numerically determined) position of both the local minimum and maximum are shifted with change of $C_0$.

In addition to the lateral tension, one may also want to induce a pressure $P$ in the lumen between the two membranes, so that $E_{tot} = E_B + E_\tau + E_P$. For a pressure close to equilibrium pressure, assuming only small volume changes, based on $dE = -PdV$ one can simplify the change in energy to $E_P = -PV$, where P is the osmotic pressure difference. The volume around the neck will be $\Delta V = A_p d - \frac{\pi}{4}\left(D_p + d\right)^2 d + \left(\frac{\pi^2}{8}d^2\left(D_p + d\right) - \frac{4\pi}{24}d^3\right)$. Then

$$\widetilde{E_{with\ P}} = \tilde{E} - \frac{PD_p^3}{\kappa}\left[\frac{s^3(8/3-\pi)}{32}+\frac{s^2\pi}{32}\right] - \frac{PD_p s}{\kappa}\frac{A_0}{8\pi} \text{ (14)}$$

$$= \tilde{f}(s) - \gamma\left[\frac{1}{16}+\frac{s(2-\pi)}{16}+\frac{s^2(3-\pi)}{16}\right] - \frac{\tilde{P}\gamma^{\frac{3}{2}}}{32}\left[s^3\left(\frac{8}{3}-\pi\right)+s^2\pi\right] - \frac{\tilde{A}\tilde{P}}{8\pi}\gamma^{\frac{1}{2}}s \text{ (15)}$$

with $\tilde{P} = P\left(\frac{\kappa}{\tau^3}\right)^{1/2}$ and $\tilde{A} = A_0\frac{\tau}{\kappa}$. Suppl. Fig. *10* shows the results of numerical minimization and how the constriction and dilation regions are retained, but similarly to spontaneous curvature a new local minimum emerges at small $\gamma$ values. The maximum's position is also shifted, however to larger $\gamma$ for higher pressure differences, increasing the region of positive slope that represents constriction.

Supplementary Note 4

#### Theory: Toroidal membrane neck; constant projected area

For constant project area, to keep control of the total surface area as well, we write our total energy as:

$$E_{tot} = E_B + E_A = E_B + \frac{K_A}{2}(A - A_0)^2 \quad (16)\text{, with}$$

$$A = 2A_p - \pi\frac{{D_p}^2}{2} - \frac{D_p d}{2}(2\pi - \pi^2) - \frac{d^2}{2}(3\pi - \pi^2) \quad (17),$$

where $E_B = f\left(\frac{D_p}{d}\right) = 4\pi\kappa\frac{D_p}{d}\frac{\left(1+\frac{d}{D_p}\right)^2}{\sqrt{1+\frac{2d}{D_p}}}tan^{-1}\left(\sqrt{\frac{2d}{D_p}+1}\right)$, $A_0$ is the targeted total surface area and a constant contribution from tension is neglected.

We define $\alpha = \frac{D_p}{d}$ and then minimise equation 8 with respect to the two diameters. Applying the necessary criterium for an optimum, $0 = \frac{\partial E_{tot}}{\partial D_p} \wedge 0 = \frac{\partial E_{tot}}{\partial d}$ yields:

$$0 = \frac{\partial E_{tot}}{\partial D_p} = \frac{f'(\alpha)}{d} + \frac{K_A}{2}\left(2A_p - \frac{\pi}{2}{D_p}^2 + \frac{D_p d}{2}(\pi^2 - 2\pi) + \frac{d^2}{2}(\pi^2 - 3\pi) - A_0\right)\left(-\pi D_p + \frac{d}{2}(\pi^2 - 2\pi)\right) \; (18)$$

$$0 = \frac{\partial E_{tot}}{\partial d} = \frac{-D_p f'(\alpha)}{d^2} + \frac{K_A}{2}\left(2A_p - \frac{\pi {D_p}^2}{2} + \frac{D_p d}{2}(\pi^2 - 2\pi) + \frac{d^2}{2}(\pi^2 - 3\pi) - A_0\right)\left(\frac{D_p}{2}(\pi^2 - 2\pi) + d(\pi^2 - 3\pi)\right) (19)$$

$$0 = \frac{\partial E_{tot}}{\partial d} + \frac{D_p}{d}\frac{\partial E_{tot}}{\partial D_p} = \frac{K_A}{2}\left(2A_p - \frac{\pi}{2}{D_p}^2 + \frac{D_p d}{2}(\pi^2 - 2\pi) + \frac{d^2}{2}(\pi^2 - 3\pi) - A_0\right)\left(D_p(\pi^2 - 2\pi) + d(\pi^2 - 3\pi) - \pi\frac{{D_p}^2}{d}\right) \; (20)$$

Considering that the diameters must have positive real values, the first term in the product in eq. (11), i.e. $(A - A_0)$, gives the following relationship between the two diameters:

$$d = D_p\left(\frac{1}{2}\frac{2-\pi}{\pi-3} + \sqrt{\frac{\pi^2-8}{4(\pi-3)^2} - \frac{4A_p - 2A_0}{\pi(\pi-3){D_p}^2}}\right) \quad (21).$$

This is a minimum of the second energy term, since $E_A = \frac{K_A}{2}(A - A_0)^2 \geq 0$ (and inserting eq. (21), $E_A = 0$). While the partial derivatives also lead to a second solution from the second term of the product in eq. (20), $D_p \approx 0.15d$, a quick 3D plot of our energy landscape clearly demonstrates that only the first solution is the unique minimum, as it lines the valley of minimal energy (Suppl. Fig. 11A).

Inserting eq. (21) into (18) & (19) we then need to ensure that $f'(\alpha) = 0$ (i.e. minimise $E_B = f(\alpha)$ ) at

$$\alpha^* = \left(\frac{1}{2}\frac{2-\pi}{\pi-3} + \sqrt{\frac{\pi^2-8}{4(\pi-3)^2} - \frac{4A_p - 2A_0}{\pi(\pi-3){D_p}^2}}\right)^{-1} \qquad (22).$$

This leaves us to minimise the energy with respect to $D_p$ (Suppl. Fig. 11B), which is done numerically over a range of constant projected areas, as in our simulation setup. The resulting optimal neck diameter $D_p(A_p)$ can be seen in Suppl. Fig. 11C and is used to evaluate the relationship between our simulation results and this theoretical description (see main text).

#### Supplementary Note 5

##### Benefits and limits of assuming toroidal neck shapes

We compared the choice for toroidal membrane necks in our theoretical model with the other commonly assumed neck shape, i.e. a catenoid. Fitting both models to the vertices of an exemplary equilibrated membrane neck (no explicitly applied lateral tension but fixed box size), yields:

Torus: $R_p + r = 14.544$, $r = 5.379$ with a loss of $L \approx 3475$ and optimality $O \approx 8.4 \times 10^{-8}$.

Catenoid: $C = 14.544$ with a loss of $L \approx 82610$ and optimality $O \approx 0.35$.

Fits were performed in 3D using a least squares method from scipy.optimize in python. $O$ represents a first-order optimality measure, which is given by the uniform norm of the gradient of the cost function at the solution.

Clearly, a neck embedded in two flat sheets of membrane in PBC seems to be well approximated by a toroidal shape, which is also highlighted in Suppl. Fig. 19 below. The toroidal assumption performs better than the catenoidal one. Here, a larger neck was chosen to increase the number of vertices available to fit. Therefore, we used a dilated neck equilibrated at constant box size (projected area), which is why the membrane is not tensionfree and $R_p/r \neq 0.6$ here.

(Performing the fit on a neck from a simulation without tension yielded:

Torus: $R_p + r = 7.486$, $r = 4.673$ with a loss of $L \approx 3374$ and optimality $O \approx 2.1 \times 10^{-6}$.

Catenoid: $C = 7.486$ with a loss of $L \approx 14648$ and optimality $O \approx 0.13$. Hence the theoretically predicted neck radius or diameter ratio of $R_p/r \approx 0.6$ was met here.)

However, the membrane necks often also display non axisymmetric shapes, even after equilibration, as highlighted by their black-and-white projections onto 2d during neck diameter analysis (Suppl. Fig. 20). This can for example be seen in constant $A_p$ −simulations at small (little membrane stretching) and expanded (more membrane stretching) $A_p$, as well as overdilated necks, with bending rigidity $\kappa = 10k_BT$. While at very small necks, the assymetry might be attributed to limits of the discrete triangulation in approximation circular shapes, at larger neck sizes this phenomenon still occurs regularly. It highlights the importance of employing simulations to capture more physical complexity than theoretical approaches might.

#### Supplementary Note 6

##### Membrane neck under constant projected area: Energy slopes (Sim. vs. Theory)

Linear fits were performed on the approximately linear part of bending energy determined in simulations for the region, where $A_p > \frac{A_0}{2}$. The linear function

$$f\left(x = \frac{2A_p}{A_0}\right) = ax + b \quad (23)$$

was fitted to the last 24 datapoints at which simulations had been performed, corresponding to the largest 24 examined $\frac{2A_p}{A_0}$. For the theoretical results predicting bending energy (see also Suppl. Fig. 11C) the ratio of slopes should correspond to the ratio of bending rigidities, if the behaviour can be well approximated linearly:

$$a_{theo}\big|_{\kappa=5k_BT} : a_{theo}\big|_{\kappa=10k_BT} : a_{theo}\big|_{\kappa=20k_BT} = 5k_BT : 10k_BT : 20k_BT = 1:2:4$$

The following results were obtained from the fit to simulation data and converted to compare the ratio of slopes between theory and simulation:

$$a_{sim}\big|_{\kappa=5k_BT} = 189.3 \pm 1.7$$

$$a_{sim}\big|_{\kappa=10k_BT} = 380.3 \pm 2.4$$

$$a_{sim}\big|_{\kappa=20k_BT} = 762.1 \pm 4.5$$

Thereby, $a_{sim}\big|_{\kappa=5k_BT} : a_{sim}\big|_{\kappa=10k_BT} : a_{sim}\big|_{\kappa=20k_BT} = 1 : 2.01 : 4.03$.

The complexity of simulation compared to theory does not cause significant deviations in the general near-linear behaviour. According to our theoretical model the near-linear behaviour should stem from the geometrical constraints of the system. The agreement in slopes together with the qualitative similarities, implies that there are no significant additional physics at play in the actual simulated system. The simulations too seem to be mainly controlled by the geometrical constraints, while the neck would otherwise dilate infinitely.

### Supplementary Figures

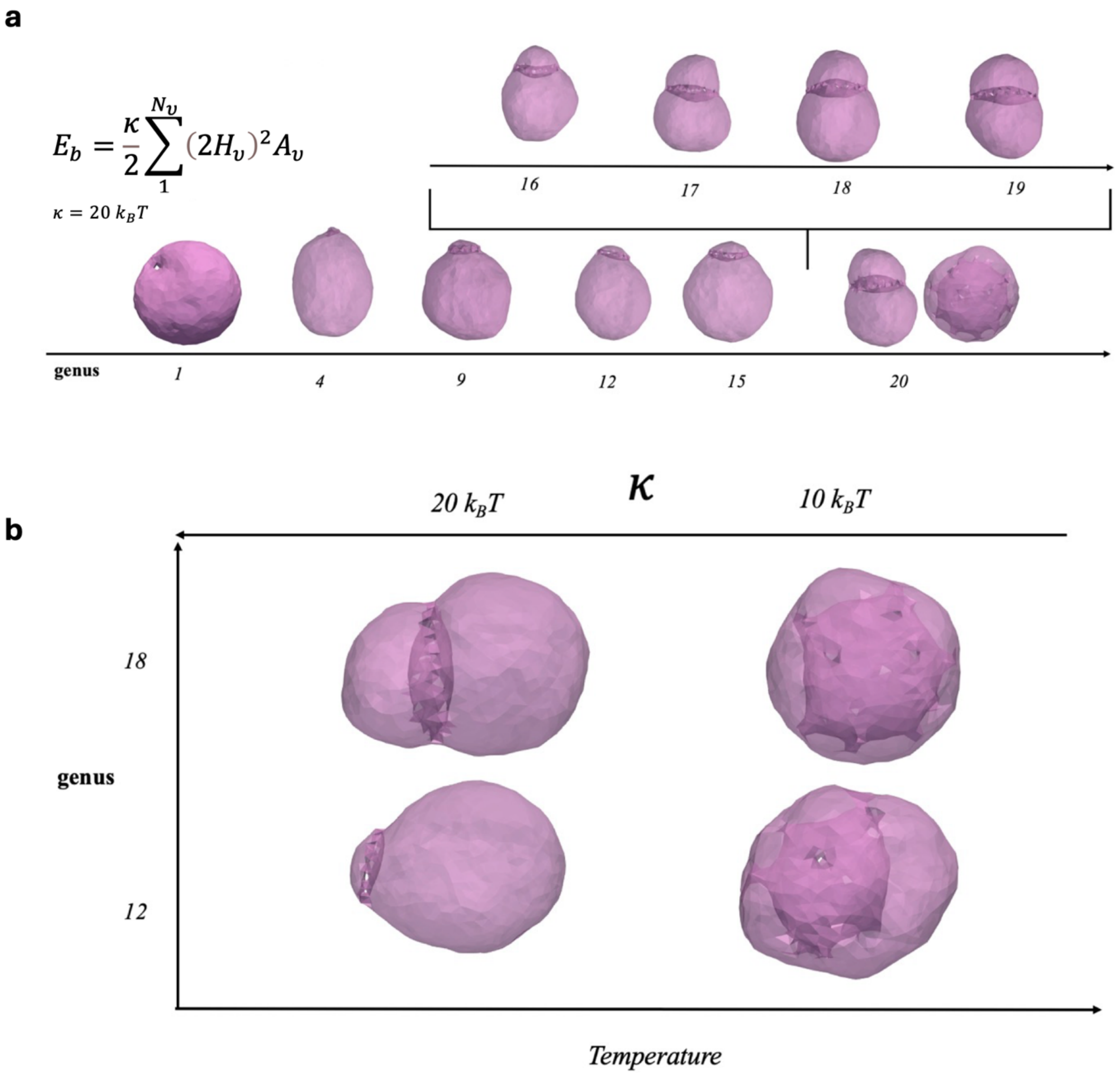

Suppl. Fig. 1 **Shapes of high-genus vesicles found after equilibration using only bending energy**. **(a)** Over a wide range of lower genera the membrane necks concentrate in one region of the vesicle. For a genus >16, they also form circular cages, while at very high-genus stomatocytes can be found. **(b)** Circular cage structures occur mainly at high bending rigidity (low temperature), where necks concentrate around a disc that divides the inner space into two compartments, whereas lower bending rigidity (high temperature) promotes spherical stomatocytes.

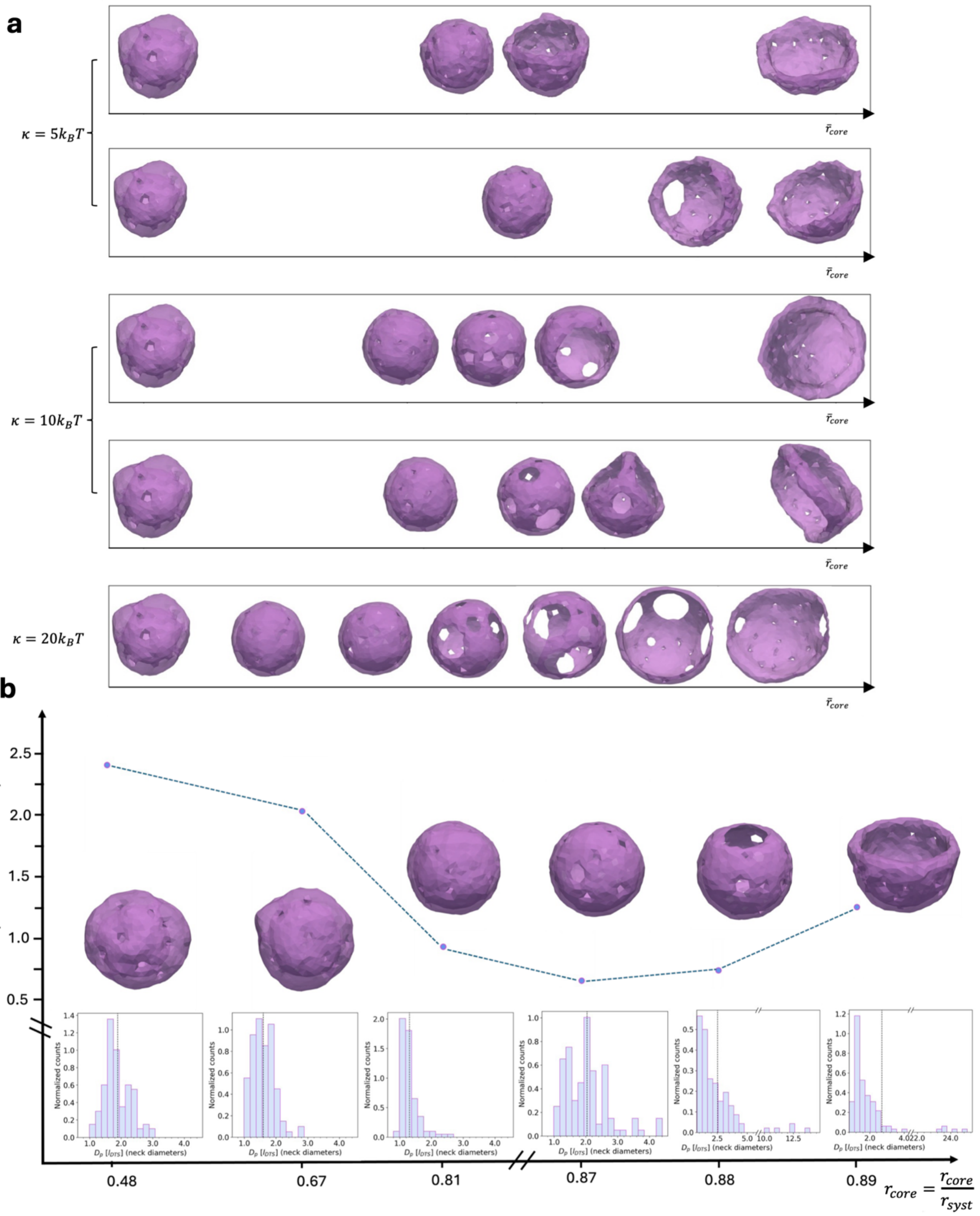

Suppl. Fig. 2 **Expanding a core bead inside a stomatocyte (g=20) leads to neck constriction, followed by eventual dilation. (a)** Changes of morphogoly during core expansion. **(b)** Evolution of neck diameters and IM-OM distance at $\kappa = 10\ k_B T$ on non-equilibrated frames.

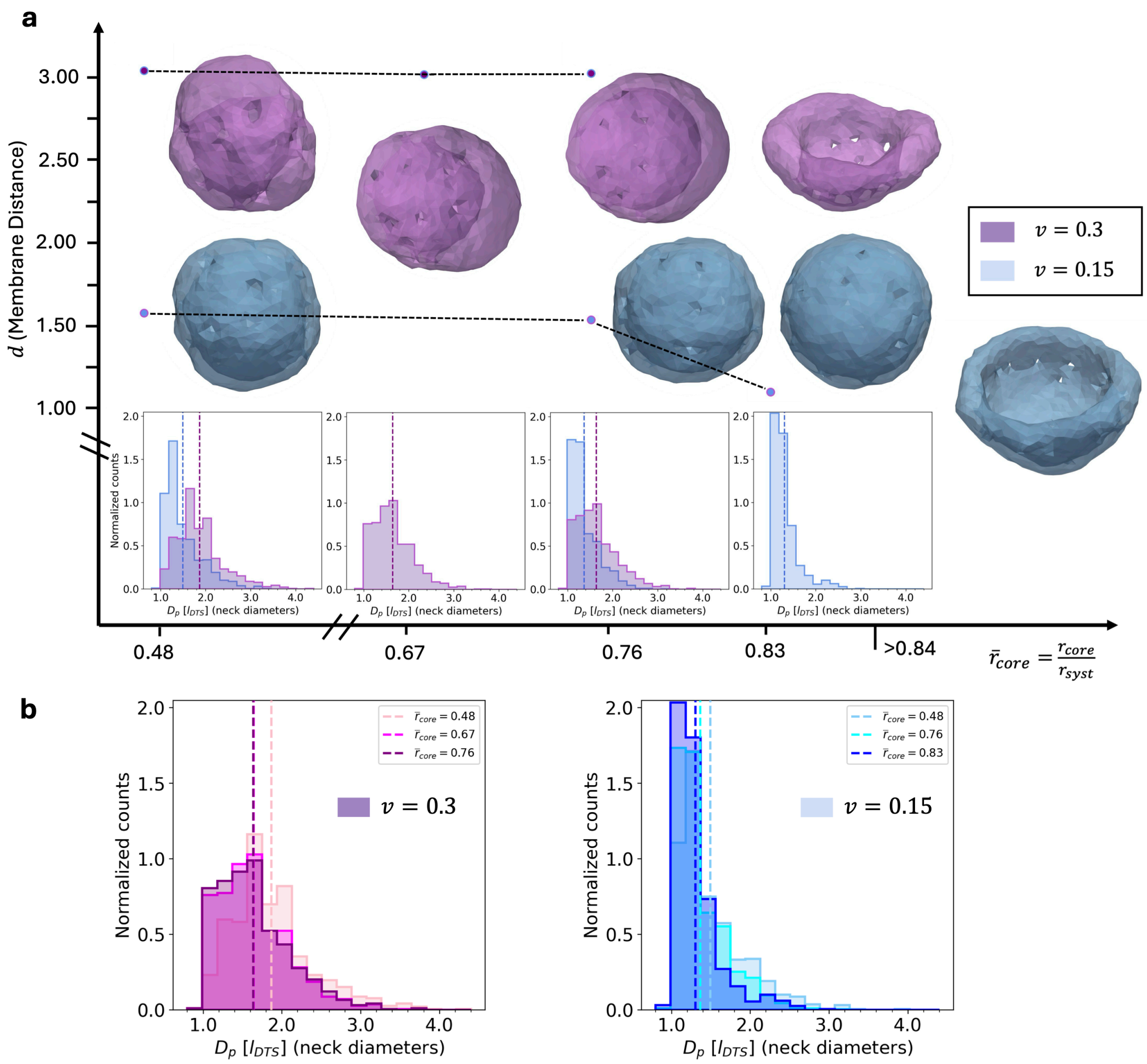

Suppl. Fig. 3 **Simulations of the genus-20 stomatocyte (21 necks) at $\kappa = 10k_BT$ under lumen volume constraint with a continuously growing core bead to represent rising osmotic pressure in the innermost compartment**. **(a)** For all spherical stomatocyte shapes (i.e. before overdilation) distributions of the neck diameters ($D_p$) are shown below the respective simulation snapshots. They were obtained from the last 50 frames of subsequent equilibrium simulations at constant core radius. The IM-OM distance ($d$) is given as the mean the same 50 frames of the average $d$ (over the membrane); errors are not shown as they are smaller than the datapoint markers. When the volume of the lumen between IM and OM is constrained, the characteristic constriction response is significantly superimposed by this effective pressure in Compartment II in addition to the core bead (Compartment I). **(b)** Direct comparison of $D_p$-distributions with growing core radius. While the constriction is still detectable in the neck diameter distributions, the neck diameters are mostly controlled by the volume of the lumen now, instead of the pressure in the innermost compartment. This highlights, how pressure in the lumen can act equivalently to membrane tension on the NE, and that controlling both the volume of the lumen and the innermost compartment may not be expedient to study the membrane neck response to increased tension, as the effects superimpose. Instead either the lumen or the innermost compartment should be constrained in order to induce clearly traceable pressure that translates to tension on the membrane. While decreasing reduced lumen volume leads to neck constriction, the same is achieved by increasing core bead radius, i.e. increasing the volume of the innermost compartment.

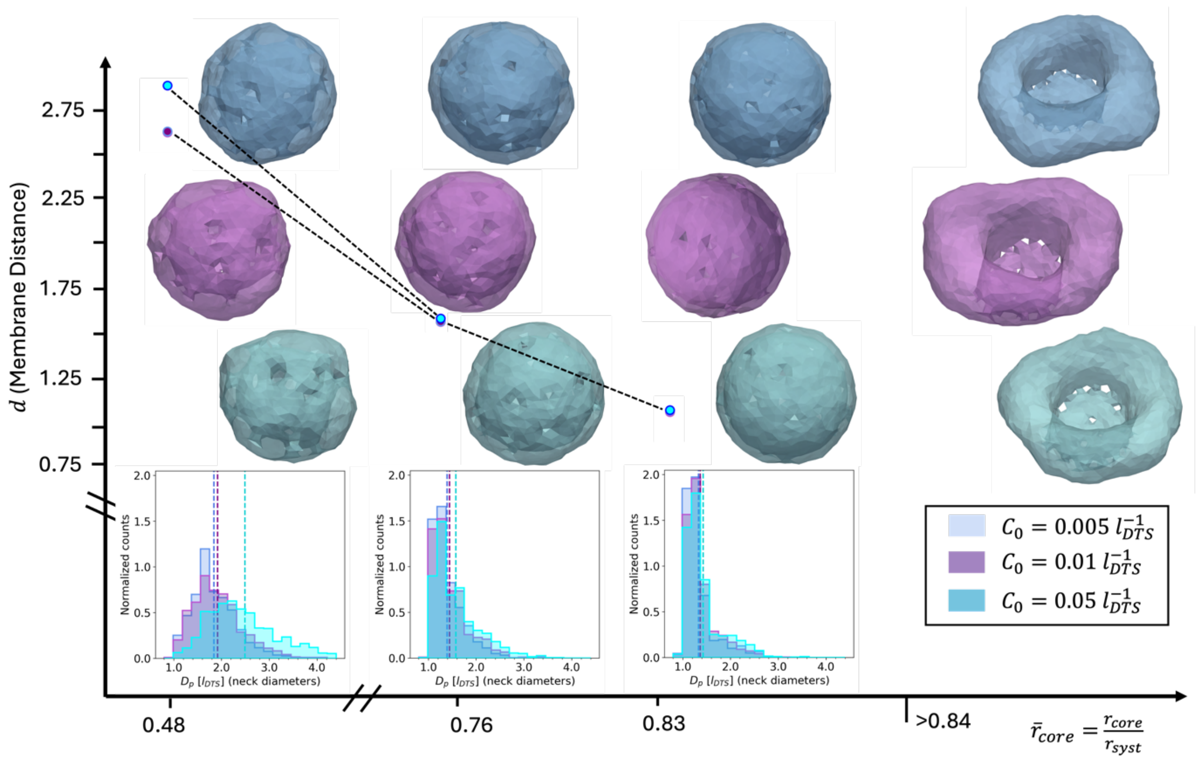

Suppl. Fig. 4 **Simulations of the genus-20 stomatocyte (21 necks) at $\kappa = 10k_BT$ with spontaneous curvature with a continuously growing core bead to represent rising osmotic pressure in the innermost compartment**. For all spherical stomatocyte shapes ($\bar{r}_{core} < 0.84$, i.e. before overdilation) distributions of the neck diameters ($D_p$) are shown below the respective simulation snapshots. They were obtained from the last 50 frames of subsequent equilibrium simulations at constant core radius. The IM-OM distance ($d$) is given as the mean the same 50 frames of the average $d$ (over the membrane); errors are not shown as they are smaller than the datapoint markers.

Clearly the characteristic constriction response followed by overdilation is maintained when spontaneous curvature is introduced. Is it slightly modified to larger neck diameters for higher spontaneous curvatures.

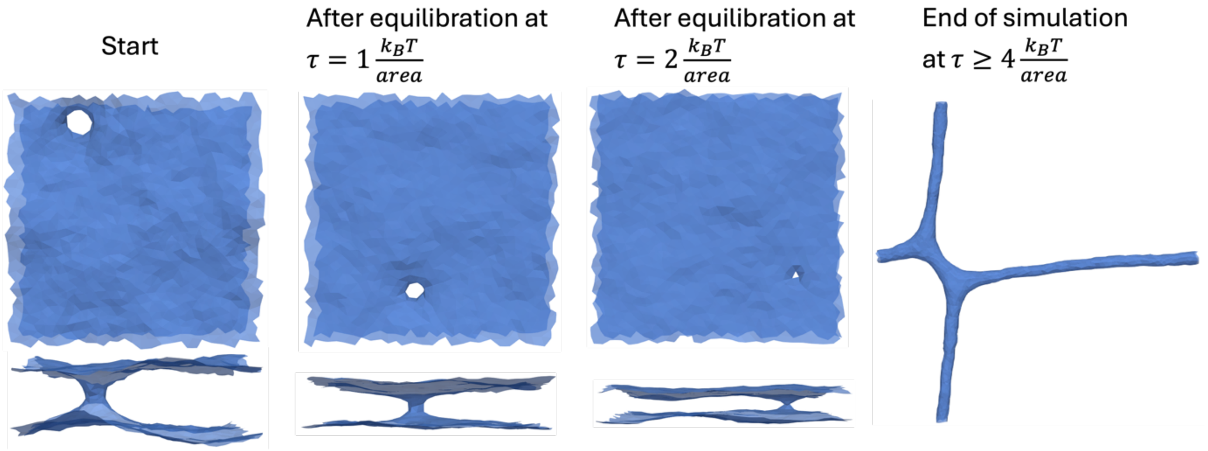

Suppl. Fig. 5 **Tension dependent neck constriction and dilation.** Membranes with bending rigidity of $\kappa = 20\ \mathrm{k_BT}$ and a starting neck diameter of $\mathrm{D_{start}} \approx 8\ \mathrm{l_{DTS}}$. Using the purified NPC diameter $\mathrm{D_p} = 43\ \mathrm{nm}$ [4,5] the lowest tensions causing dilation in simulations (see results 1.) convert to physical units as $\tau_D|_{\kappa=5k_BT} \approx 1.5 \times 10^{-4}\frac{N}{m}$, $\tau_D|_{\kappa=10k_BT} \approx 2.2 \times 10^{-4}\frac{N}{m}$ and $\tau_D|_{\kappa=20k_BT} \approx 5.9 \times 10^{-4}\frac{N}{m}$, by comparing the simulated $\mathrm{D_{p,sim}} \approx 8\mathrm{l_{DTS}} =$ 43nm giving $\mathrm{l_{DTS}} \approx$ 5.4nm (see also Methods). In good agreement, assuming $\mathrm{D_p} = 43\ \mathrm{nm}$ as the neck diameter, our theoretical model (Supplementary Note 3) predicts $\tau_{crit}|_{\kappa=5k_BT} \approx 1.2 \times 10^{-4}\frac{N}{m}$, $\tau_{crit}|_{\kappa=10k_BT} \approx 2.5 \times 10^{-4}\frac{N}{m}$ and $\tau_{crit}|_{\kappa=20k_BT} \approx 4.9 \times 10^{-4}\frac{N}{m}$.

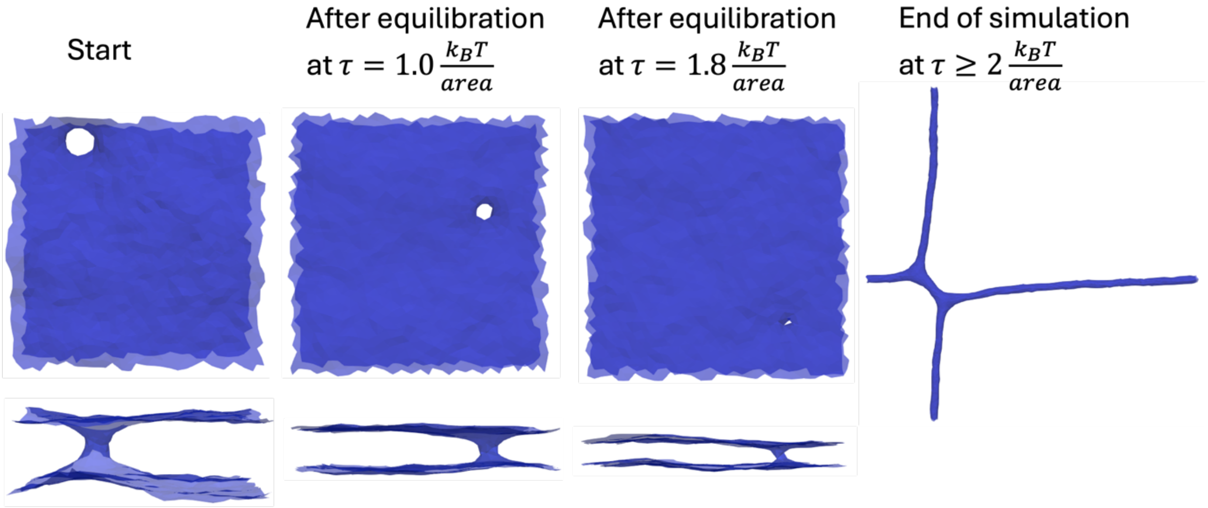

Suppl. Fig. 6 **Tension dependent neck constriction and dilation with spontaneous curvature $C_0 = 0.01\ l_{DTS}^{-1}$.** Membranes with bending rigidity of $\kappa = 20\ k_BT$ and a starting neck diameter of $D_{start} \approx 8\ l_{DTS}$.

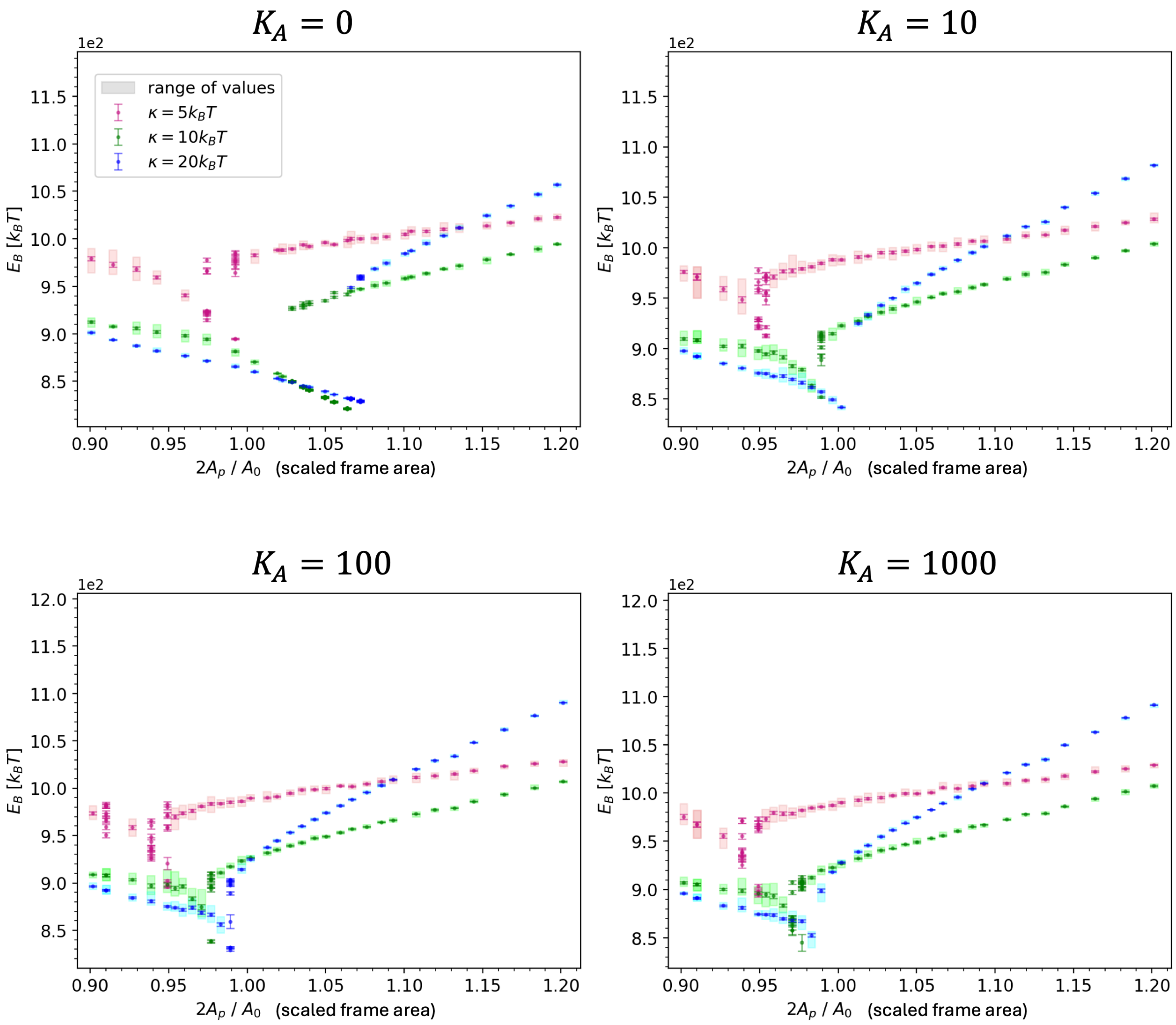

Suppl. Fig. 7 **Effect of area compressibility in simulations over constant projected area.** Average bending energy reached after equilibration in systems with a membrane neck under PBC over a range of projected areas. Each datapoint represents the mean energy averaged over 10 simulation replicas with the standard error of the mean. At branching points all replicas are displayed instead of the mean. An error on each replica was obtained by block averaging over the last $10^6$MC steps (20 blocks, length of 500 energy outputs; energy was saved every 100 MC steps). Increasing area compressibility for fixed reference area $A_0 = 2557\ l_{DTS}^2$, moves the transition point to projected areas $2A_p < A_0$, as less surface area can be gained by stretching. Therefore, the less stretchable the surface area is, the earlier the system needs to dilate necks to increase the projected area further.

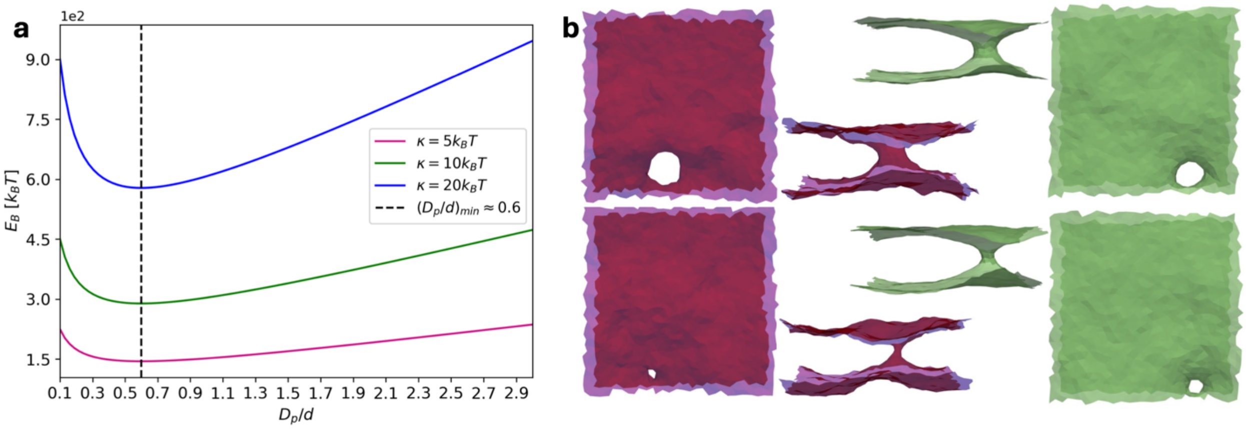

Suppl. Fig. 8 **(a) Bending energy of a toroidal membrane neck as a function of the two characterising torus diameters for different bending rigidities. (b) The degeneracy of the neck size in the tensionless setup.** The snapshots are taken from different points in the simulation after equilibration was reached showing some of the maximal and minimal neck sizes between which the equilibrium configurations fluctuate; (left/ green) bending rigidity of $\kappa = 10k_BT$ ; (right/red) $\kappa = 5k_BT$. The examples highlight that in the tensionless environment neck diameters can vary significantly, although equilibration is successful with similar final bending energies being reached in all replicas.

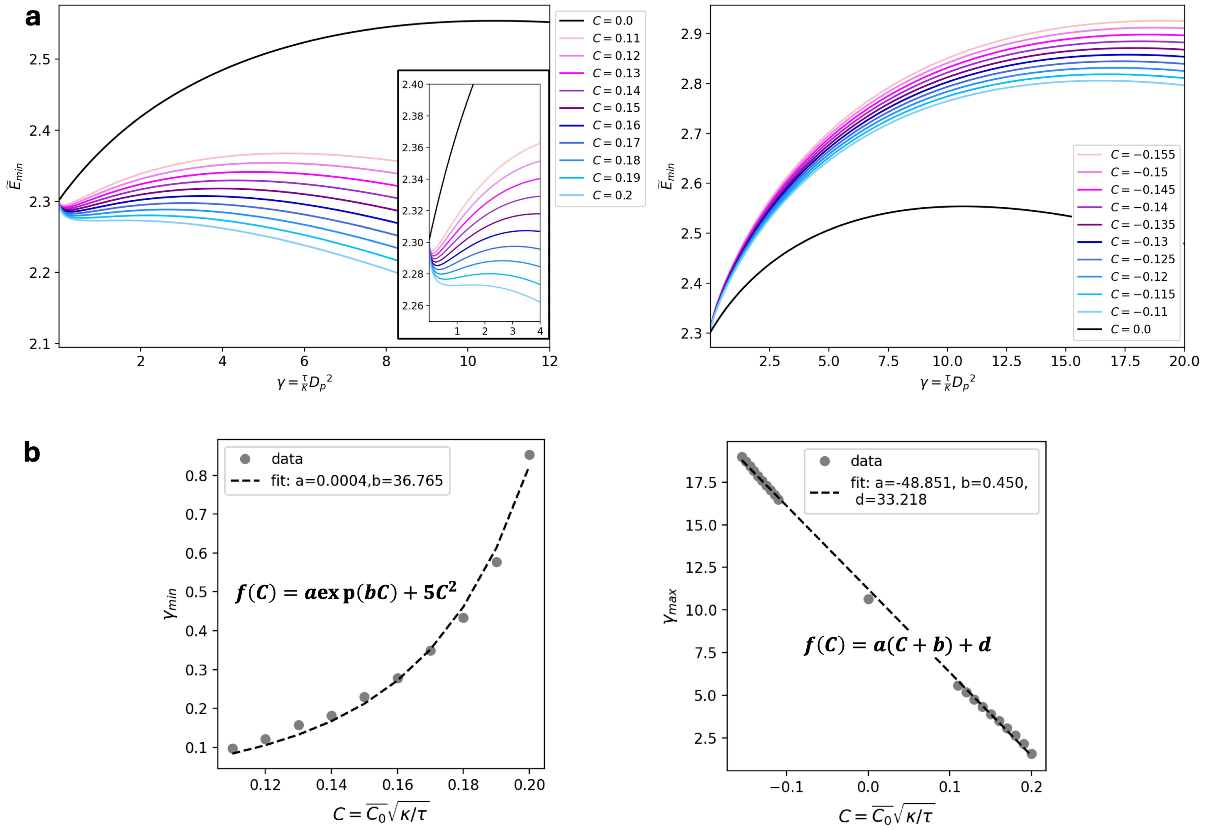

Suppl. Fig. 9 **(a) Minimal rescaled total energy of a toroidal membrane neck with spontaneous curvature $C_0$ under tension. (b) Positions of local maximum $\gamma_{max}$ and minimum $\gamma_{min}$, if present, as a function of spontaneous curvature.** Higher spontaneous curvatures, at a given tension & bending rigidity, shift the maximum to smaller neck diameters until the constriction regime disappears. The minimum is only present for a certain range of spontaneous curvatures and is shifted to larger neck diameters with larger $C = \sqrt{\kappa \backslash \tau}\, C_0$ . The fits quantify this relationship in more detail.

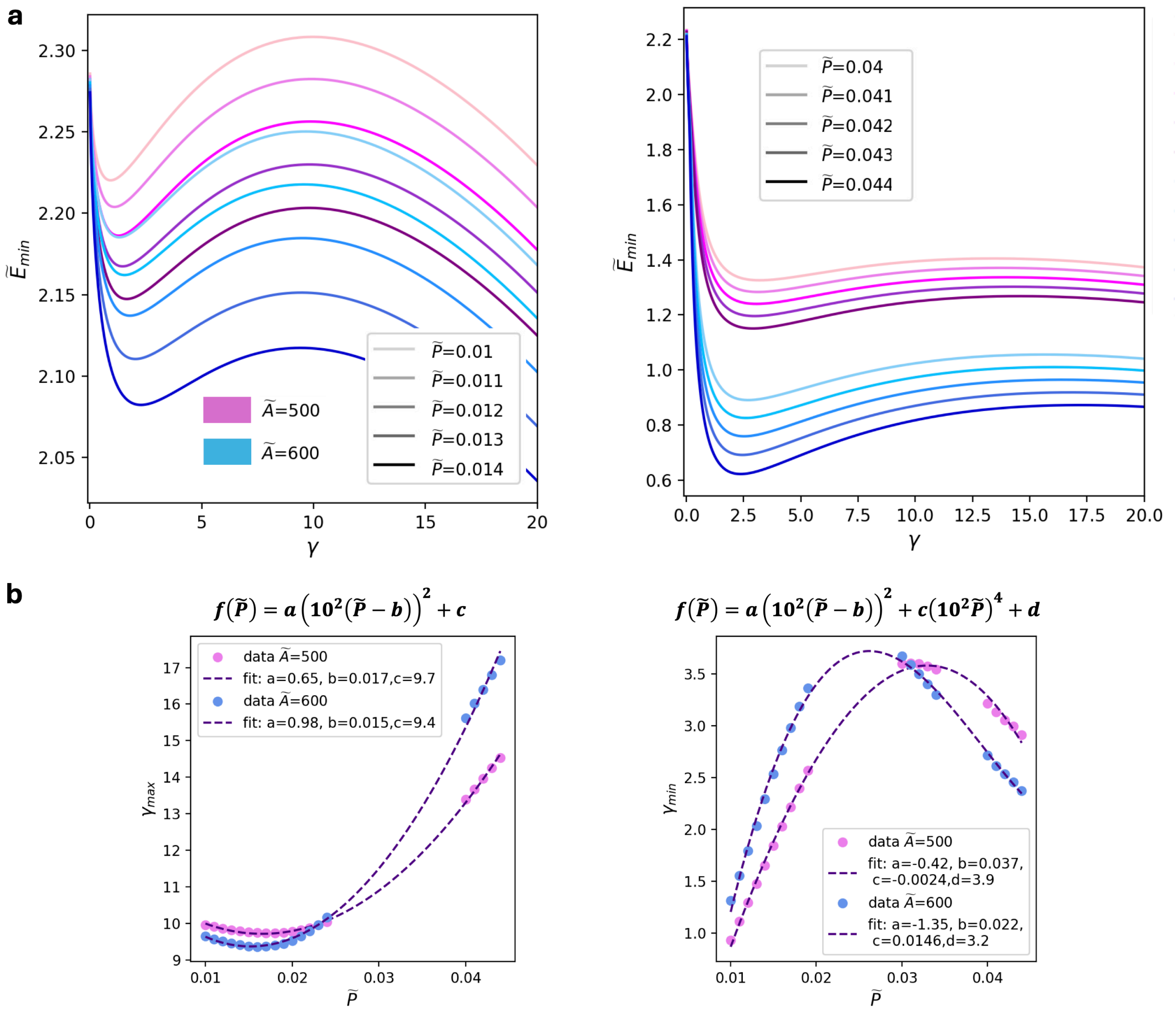

Suppl. Fig. 10 **(a) Minimal rescaled total energy of a toroidal membrane neck under tension and pressure in the lumen. (b) Positions of local maximum $\gamma_{max}$ and minimum $\gamma_{min}$, as a function of rescaled pressure.**

Here, $\gamma = \frac{\kappa}{\tau} D_p^2$, $\tilde{P} = P\left(\frac{\kappa}{\tau^3}\right)^{\frac{1}{2}}$and $\tilde{A} = A_0 \frac{\tau}{\kappa}$, where $A_0$ is the surface area of the membrane and $P$ the (osmotic) pressure difference between lumen and environment. Higher pressure, at a given tension & bending rigidity, shifts the maximum to larger neck diameters favoring constriction and delaying dilation. The minimum is shifted to smaller neck diameters with larger $\tilde{P}$. For very small pressure differences the relationships inverse. The fits quantify the relationship in more detail.

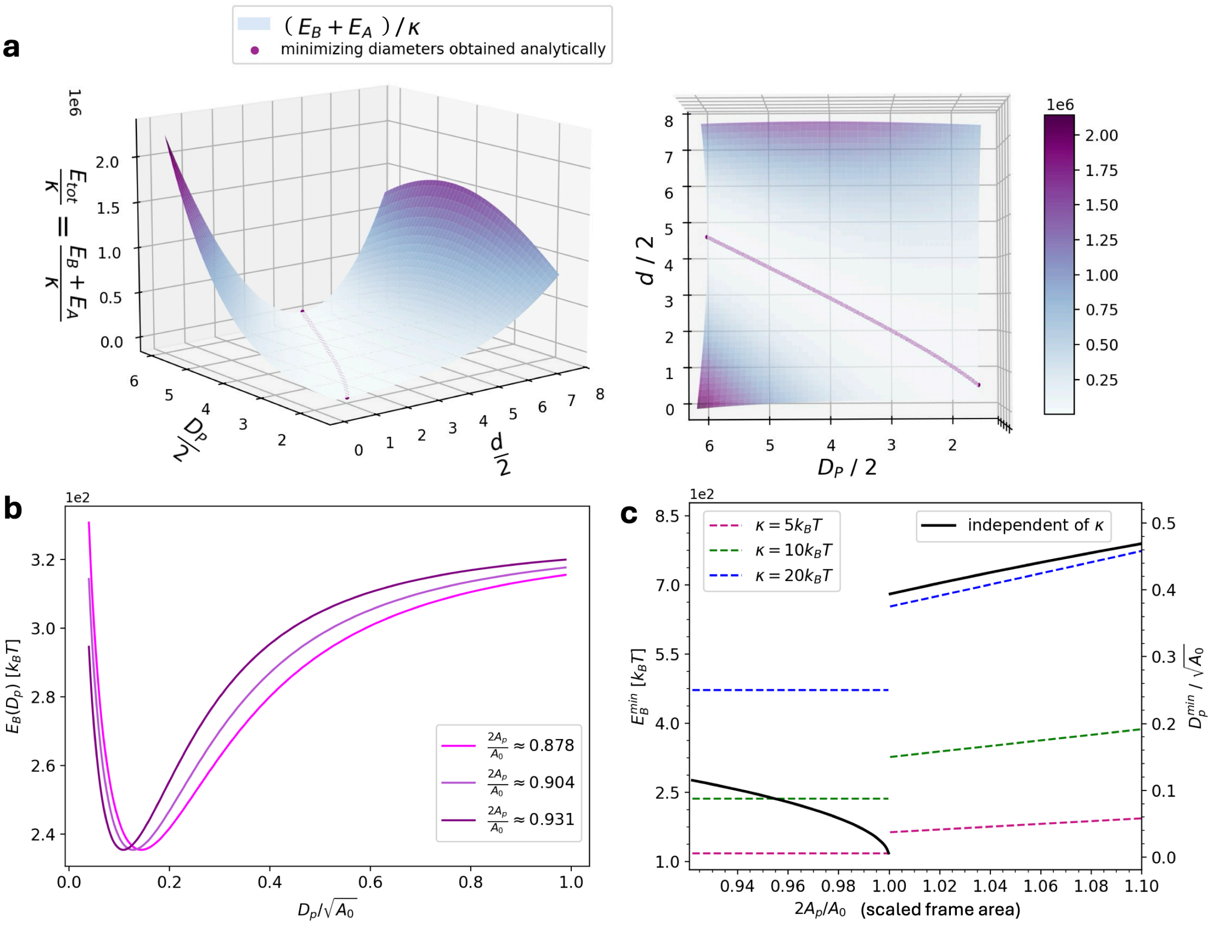

Suppl. Fig. 11 **Theoretical evaluation of a toroidal membrane neck in PBC assuming constant projected area and adding a potential for surface area control.**

**(a)** Total energy landscape. Since lateral tension constitutes only a constant term due to the fixed projected area, it is neglected here. The dotted purple line showes the path of the analytically obtained relationship between diameters $D_p, d$ that minimises the energy.

**(b)** Total energy as a function of neck diameter $D_p$ using the minimising relationship $d(D_p)$ at three different values of projected area. $D_p$ has been scaled with the target area for generalisation to dimensionless variables. The target area was chosen to be similar to surface areas in simulations. Since the minimisation leads to the area control term of the total energy being zero, and we neglegt the constant tension contribution, only bending energy remains.

**(c)** Results of minimising the energy for the remaining variable $D_p$. The left both the reached minimal energy, and the minimising diameters are shown. The results for diameters are independent of bending rigidity $\kappa$, but scale with the targeted surface area $A_0$.

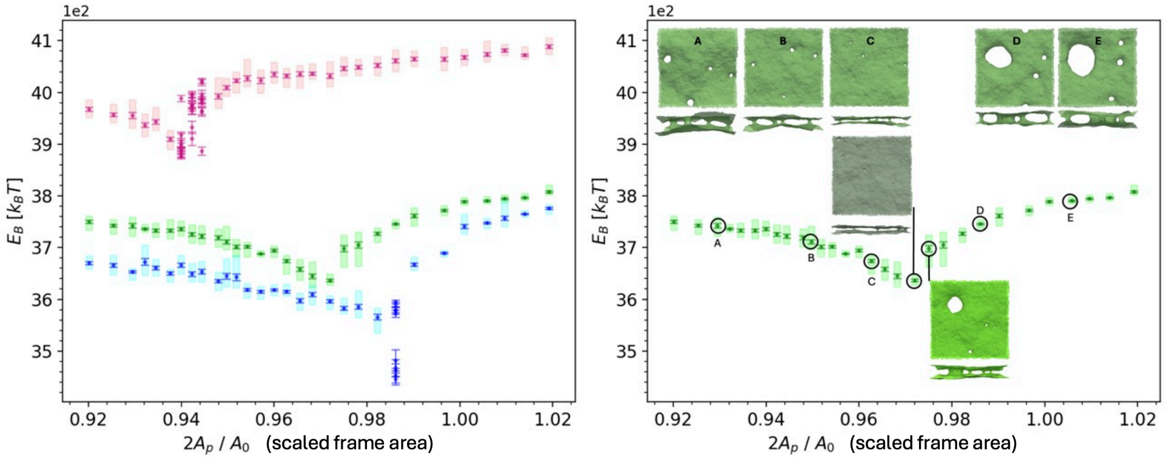

Suppl. Fig. 12 **Four-neck system in PBC: Bending energy as a function of projected area for different bending rigidity.** Results were obtained from simulation with near-constant surface area $A_0 = 10936\, l_{DTS}^2$ ($K_A = 1000 k_B T$). For each data point, 10 replica simulations were performed. Each point corresponds to their mean energy with the standard error of the mean. At transition points all replicas are displayed instead of the mean. An error on each replica was obtained by block averaging. Similar to the system with two necks, the four necks system also only exhibits one transition point, after which one of the necks dilates continously, while the others stay relatively constricted. This is visualised for the case of $\kappa = 10 k_B T$ with representative snapshots from simulations after equilibration.

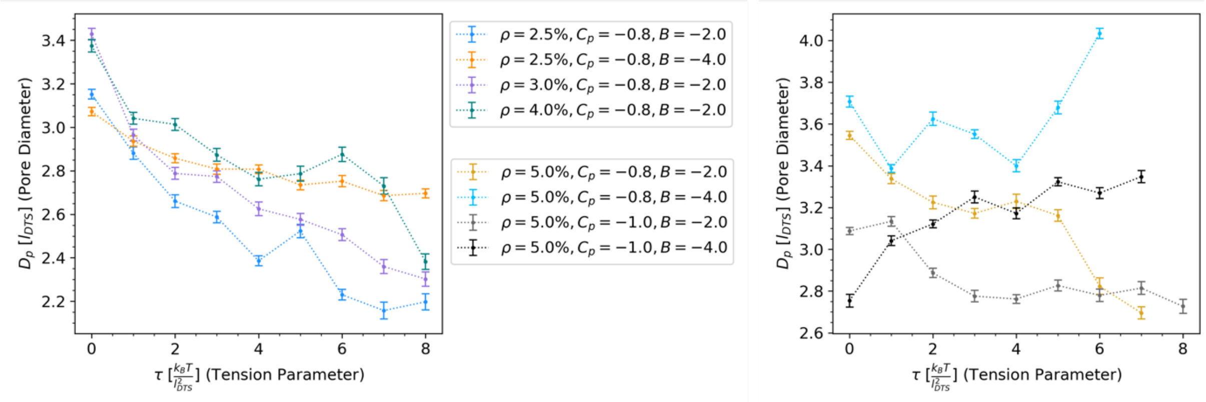

Suppl. Fig. 13 **Diameter response to tension of a membrane neck in PBC decorated by proteins.** Diameter values were obtained as the mean of the 50 last outputs from equilibrated simulations (output every 2000 MC steps), with standard error of the mean. They are shown over applied tension parameter in simulations for different protein abundance ($\rho$ denotes the percentage of membrane vertices with an inclusion, i.e. a protein), curvature preference ($C_p = C_\parallel$in $[l_{DTS}^{-1}]$) and interaction strength ($A = 1 k_B T, B$ as specified in legend).

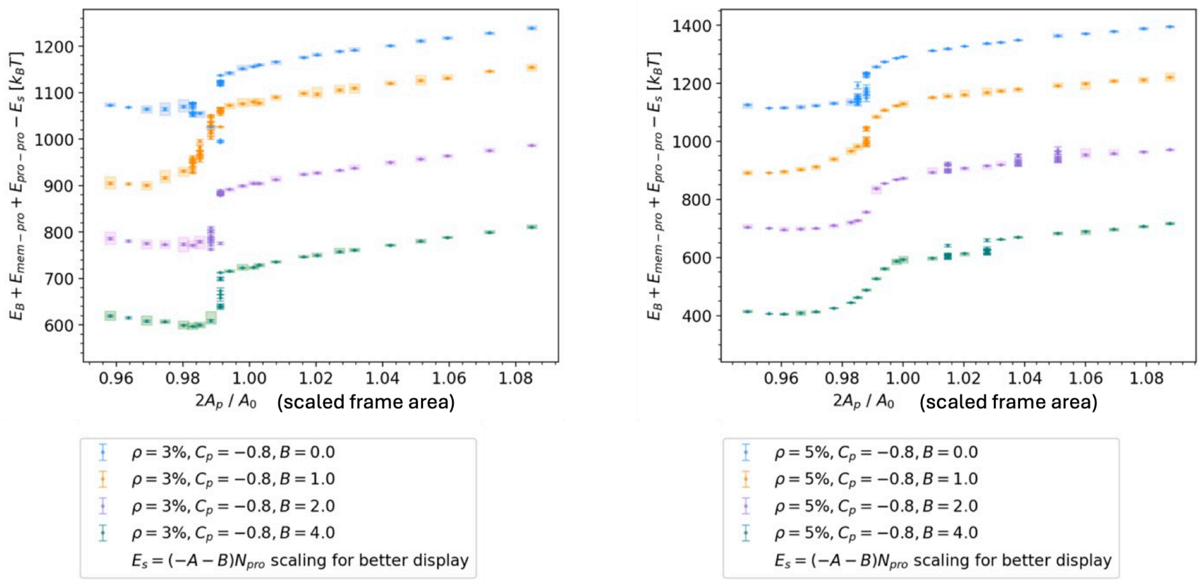

Suppl. Fig. 14 **Constant box (constant projected area) simulations of double membranes connected by a neck with protein assembly in PBC.**
Average equilibrium bending energies and local energies induced by proteins over a range of constant projected areas, for different protein complex sizes (left vs right, $\rho$ denotes the percentage of membrane vertices with an inclusion, i.e. a protein). The protein interaction strengths were varied ($A = 1k_BT, B$ as specified in legend) at constant curvature preference ($C_p = C_{\parallel}$in $[l_{DTS}^{-1}]$). Results were obtained from simulations with near-constant surface area ($K_A = 1000\ k_BT$). Datapoints correspond to mean values of 10 replicas and errors were obtained as the standard error of the mean. Individual replicas at transition points are shown without errors.

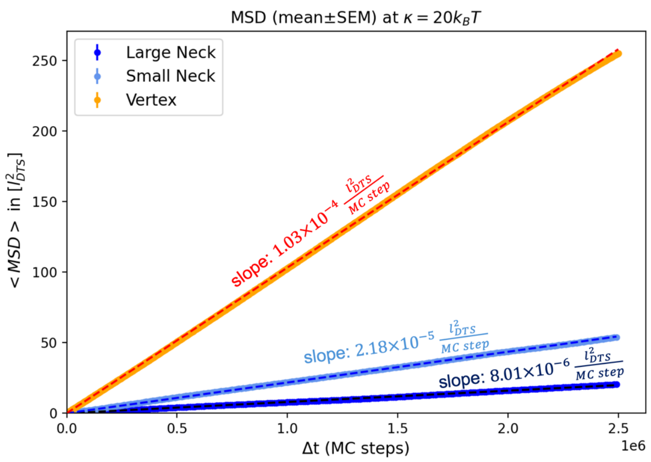

Suppl. Fig. 15 **Lateral mobility of necks in constant box (constant projected area) simulations of double membranes connected by a neck in PBC.** We analysed the average MSD over the course of simulations with 10 replica each over $10 \times 10^6$MC steps for a large ($D_p \approx 6\ l_{DTS}$) neck and a small neck ($D_p \approx 3\ l_{DTS}$) connecting double membrane patches in PBC. The diffusion is compared to that of vertex positions (≈ 2000 vertices) in flat membrane patches in PBC that were also simulated for $10 \times 10^6$MC steps. The error on the average MSD at each $\Delta t$ is the standard error of the mean of the different replica or vertices, respectively.

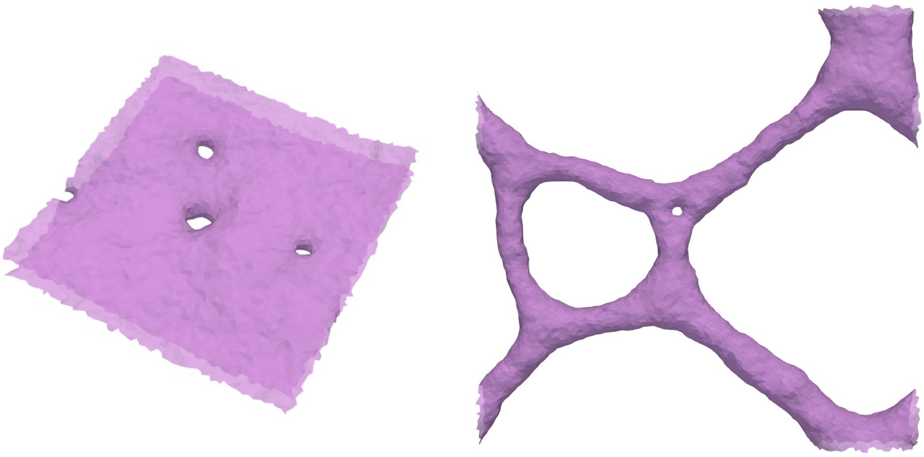

Suppl. Fig. 16 **Snapshots at the start of and during a simulation of four necks system with PBC under large lateral tension that leads to dilation**. While most necks dilate, a structure resembling the characteristic three-way junctions of the endoplasmic reticulum forms.

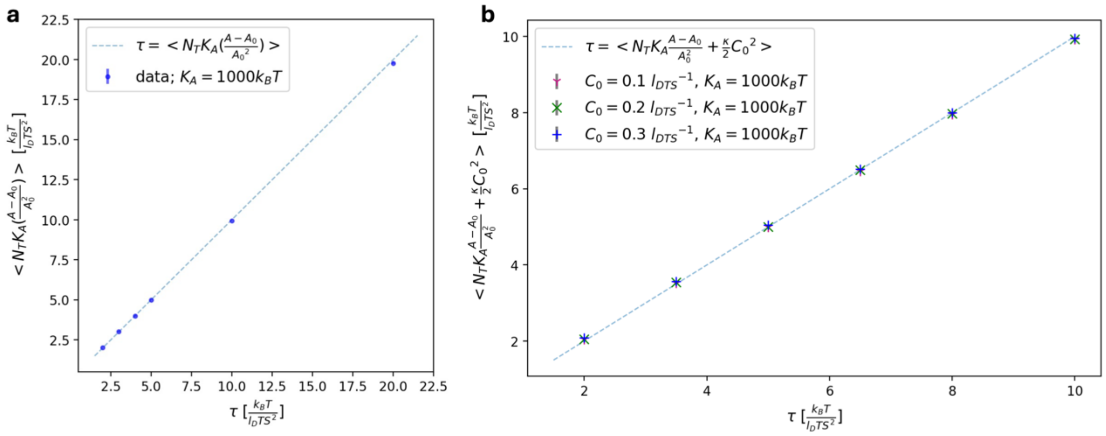

Suppl. Fig. 17 **Membrane stress balance.** Simulations were performed for $5 \times 10^6$ MC steps of which the last $10^6$ were analyzed. Errorbars (smaller than data markers) represent the standard error of the mean obtained from three replicas of simulations at each datapoint. A flat membrane segment was simulated in periodic boundary conditions in FreeDTS in the constant frame tension ensemble $(N, \tau, T)$ with additional surface area constraint according to eq. (5) (main text) with area compressibility $K_A = 1000 k_B T$.
**(a)** Average internal stresses (y-axis) versus externally imposed tension tau (x-axis). This is consistent with equation 1 of Lipowsky, Faraday Discuss (2025) [6]. **(b)** Average total tension from stretching and spontaneous tension, emerging from spontaneous curvature (y-axis) versus externally imposed tension tau (x-axis).

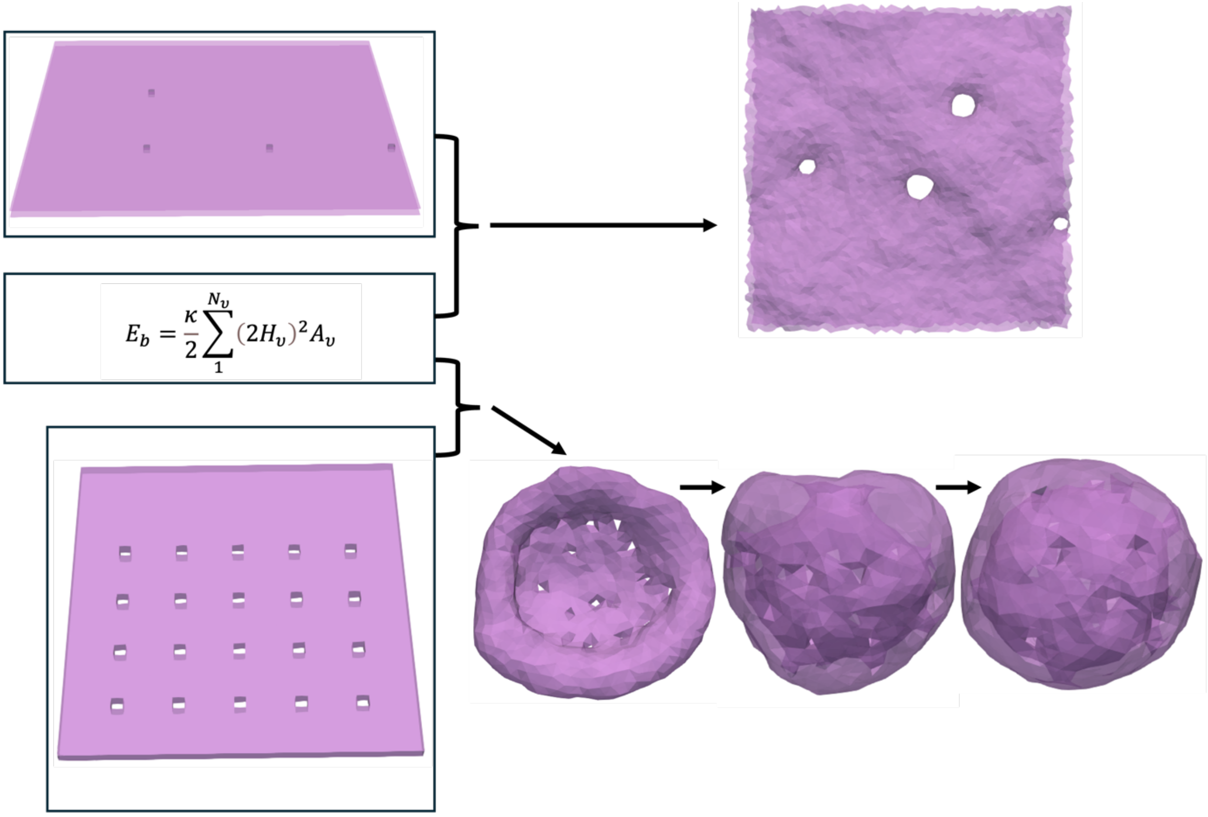

Suppl. Fig. 18 **The process of generating realistic, relaxed membrane configurations from the starting configurations**: (top), example with four membrane necks in PBC (bottom) for closed systems (g=19).

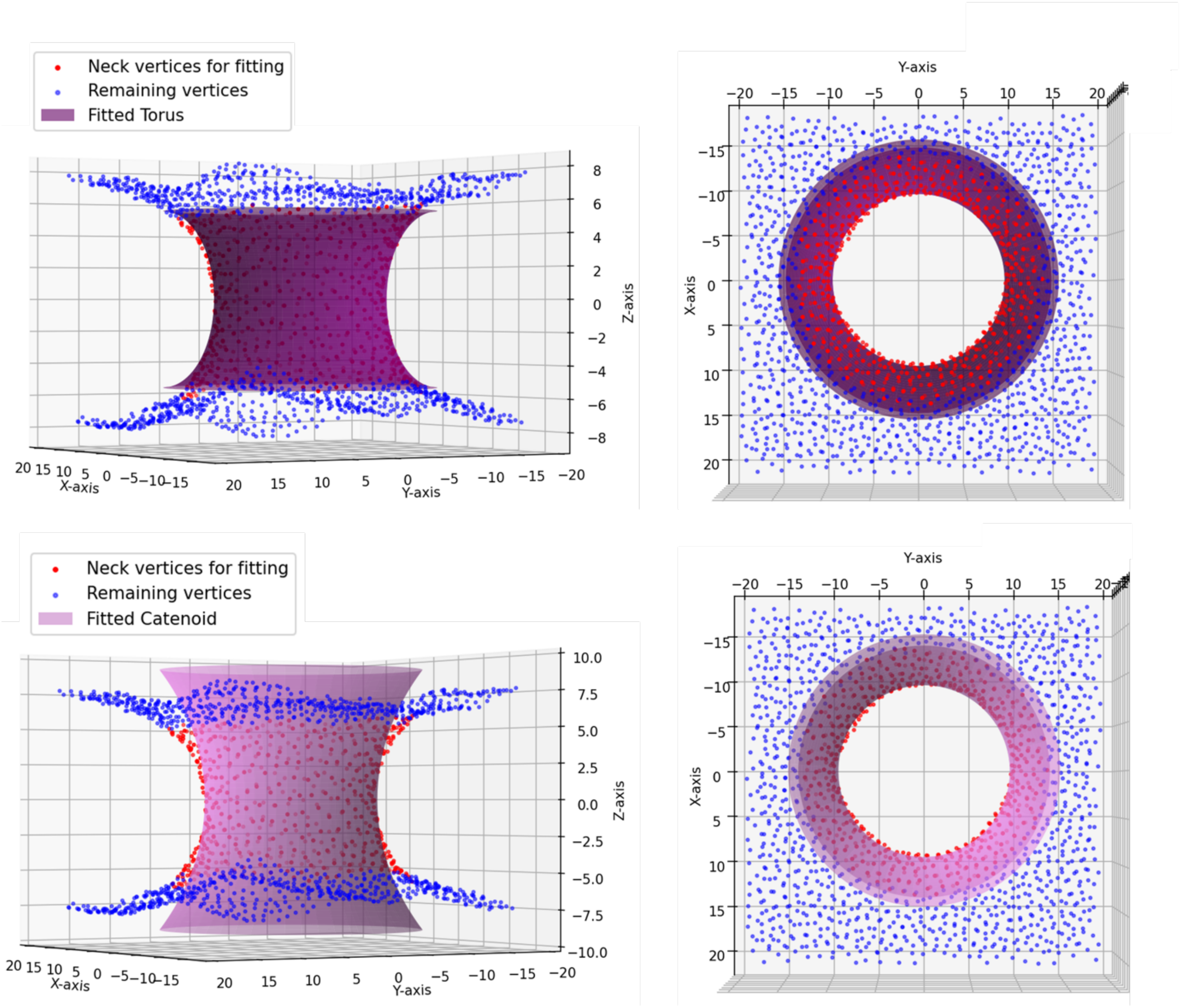

Suppl. Fig. 19 **Membrane mesh vertex data of an equilibrated neck and fitted torus (top) and catenoid (bottom).**

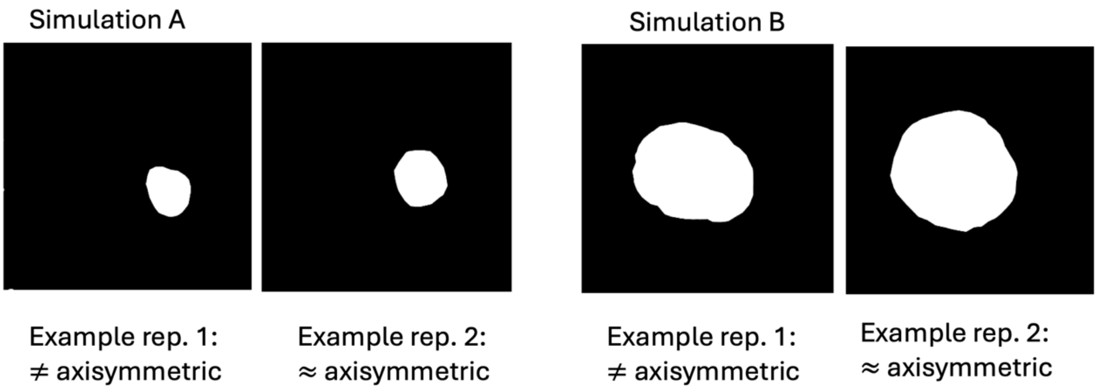

Suppl. Fig. 20 **Examples of non-axisymmetric and approximately axisymmetric shapes of membrane necks in simulations after equilibration, visualised by binary images of the membrane projected onto the x-y-plane.** Simulation A shows a smaller neck at smaller projected area, Simulation B a larger neck at bigger projected area. All replicas were equilibrated at their respective constant boxsize.

#### Supplementary References

1. Zimmerli, C. E. *et al.* Nuclear pores dilate and constrict in cellulo. *Science* **374**, eabd9776 (2021).
2. Lammerding, J. Mechanics of the Nucleus. *Compr. Physiol.* **1**, 783–807 (2011).
3. Maul, G. & Deaven, L. Quantitative determination of nuclear pore complexes in cycling cells with differing DNA content. *J. Cell Biol.* **73**, 748–760 (1977).
4. Schuller, A. P. *et al.* The cellular environment shapes the nuclear pore complex architecture. *Nature* **598**, 667–671 (2021).
5. Lin, D. H. & Hoelz, A. The Structure of the Nuclear Pore Complex (An Update). *Annu. Rev. Biochem.* **88**, 725–783 (2019).
6. Lipowsky, R. The many faces of membrane tension for biomembranes and vesicles. *Faraday Discuss.* **259**, 234–263 (2025).
